# Thermodynamic Architecture and Conformational Plasticity of GPCRs

**DOI:** 10.1101/2022.11.26.518034

**Authors:** Sathvik Anantakrishnan, Athi N. Naganathan

## Abstract

G-protein-coupled receptors (GPCRs) are ubiquitous integral membrane proteins involved in diverse cellular signaling processes and consequently serve as crucial drug targets. Here, we carry out the first large-scale ensemble thermodynamic study of 45 different ligand-free GPCRs employing a structure-based statistical mechanical framework and identify extensive conformational plasticity encompassing the seven transmembrane (TM) helices. Multiple partially structured states or intermediates co-exist in equilibrium in the native ensemble, with the TM helices 1, 6 and 7 displaying varied degrees of structure, and TM3 exhibiting the maximal stability. Active state GPCRs are characterized by reduced conformational heterogeneity with altered coupling-patterns distributed not just locally but throughout the structural scaffold. Strongly coupled residues are distributed across the structure in an anisotropic manner accounting for only 13% of the residues, highlighting that a large number of residues in GPCRs are inherently dynamic to enable structural motions critical for function. Our work thus uncovers the thermodynamic hallmarks of GPCR structure and activation, and how differences quantifiable only via higher-order coupling free energies provide insights into their exquisite structural specialization and the fluid nature of the intramolecular interaction network. The intricate landscapes and perturbation methodologies presented here lay the foundation for understanding allosteric mechanisms in GPCRs, location of structural-functional hot-spots, and effects of disease-causing mutations.

## Introduction

G protein-coupled receptors (GPCRs) are a large superfamily of integral membrane proteins found across the eukaryotic tree of life that are involved in numerous critical signaling processes. The human genome is known to contain over 800 different GPCRs with roles in vision, taste, smell, neurotransmission, immunoregulation, homeostasis, and growth.^1^ Their physiological importance and the variety of processes in which they are involved are well illustrated by the fact that over 30% of clinically-approved drugs target GPCRs.^2^ Mutations in GPCRs have been implicated in a wide variety of diseases including retinitis pigmentosa, thyroid disease, epilepsy, fertility disorders, and carcinomas.^3,4^

GPCRs are divided into six classes based on their functions and sequence homology, with the class A (Rhodopsin-like) receptors comprising the largest group. All GPCRs share a common transmembrane domain structure consisting of seven helices arranged in a highly conserved topology (Figure 1A). This transmembrane domain, also called the 7TM domain, is involved in ligand binding, allosteric signal transduction, and the binding and subsequent activation of downstream effector proteins. Ligand-binding on the extracellular side of the helix bundle induces allosteric conformational changes that result in G-protein binding and activation on the intracellular side.^5^ Signaling through GPCRs is induced by a wide variety of stimuli, including heat, mechanical stresses, small molecules, and peptides. Signals are transmitted within the cell through signaling transducers, heterotrimeric G proteins and β-arrestins. GPCRs contain several highly conserved sequence motifs and structural features. The first and second extracellular loops (ECL1 and ECL2) contain conserved cysteine residues that form a disulphide bridge. On the intracellular side, many GPCRs in their inactive state contain an “ionic lock”, a network of salt bridges between residues in TM3 and TM6 (Glu6.30, Arg3.50, and Asp3.49 in β_2_-AR using the Ballesteros–Weinstein numbering scheme).^5^ The helical bundle contains multiple conserved sequence motifs that act as “microswitches” which are involved in GPCR activation and stabilize the conformation of the transmembrane helices in the active state. These include the D[E]RY sequence in TM3, the NPxxY motif in TM7, and the PIF motif at the interface of TM3, TM5, and TM6.^6,7^

**Figure 1.**
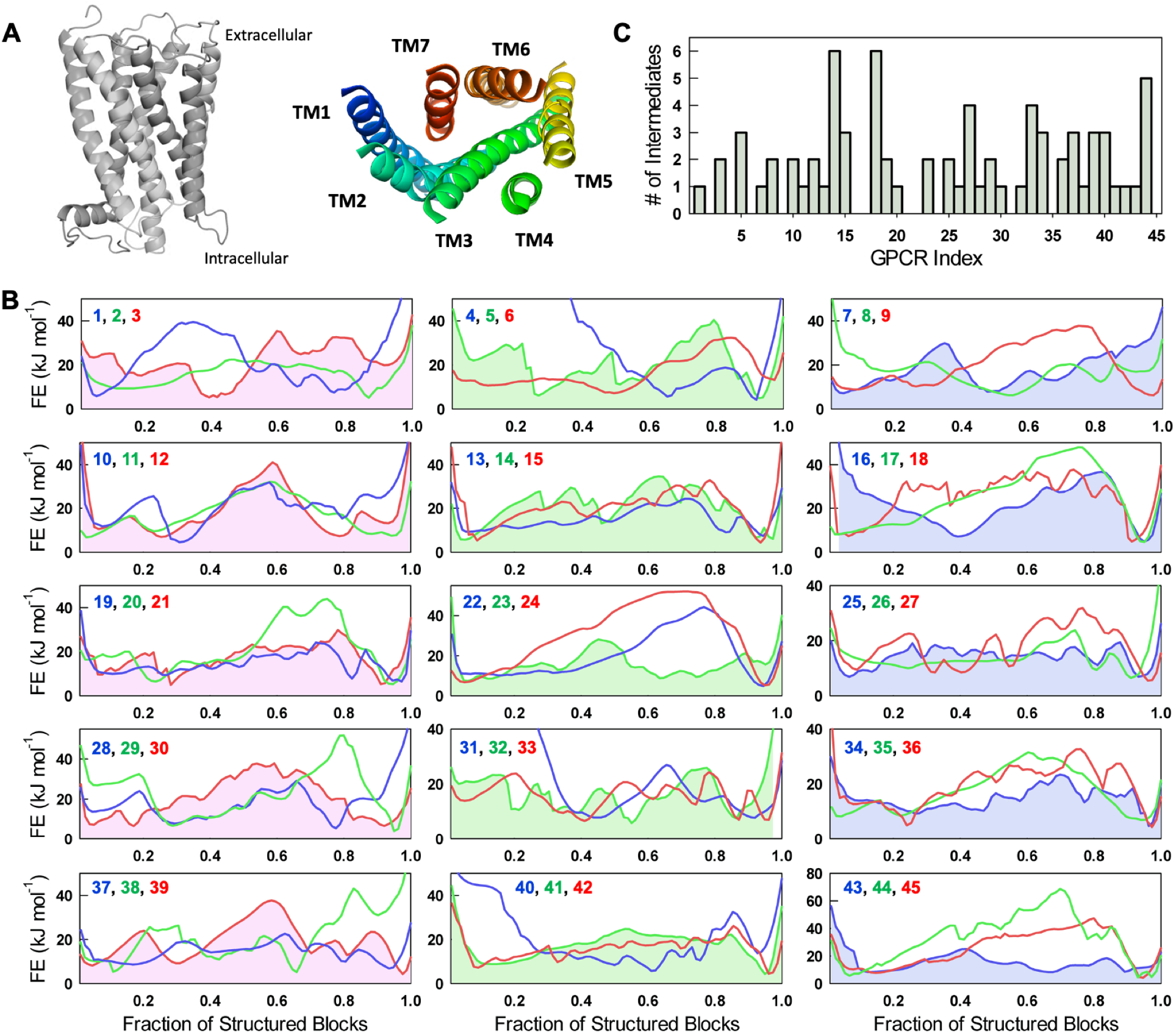
GPCR free energy profiles. (A) Representative structure of GPCR with the seven transmembrane (TM) helices viewed side on (left) and from the extracellular side (right). (B) One-dimensional free energy (FE) profiles of the 45 GPCRs studied in this work, as a function of the reaction coordinate, the fraction of structured blocks, at the melting temperature. The more folded conformations exist to the right of the free energy profile (high reaction coordinate values), while the partially structured states and unfolded conformations will be populated to the left (intermediate and low reaction coordinate values, respectively). The numbers on the top left of every panel are the GPCR indices. (C) The number of intermediates determined from the free energy profiles in panel B assuming a *1RT* threshold.

Despite the importance of GPCRs and their ubiquitous presence in eukaryotic species, the extent of native ensemble heterogeneity in GPCRs is an open question. Their presence in the membrane makes their purification and reconstitution for biophysical experiments difficult. Further, the responses of various GPCRs to inductive stimuli take place on timescales ranging from milliseconds to hours.^8^ Molecular dynamics (MD) simulations of the folding of these large receptors (the TM helices alone are ~300 residues in length) in their native membrane environment over large timescales are computationally challenging. Aside from all-atom MD simulations,^9–15^ multiple biophysical techniques have been used to probe GPCR conformational dynamics, ligandbinding, and GPCR-G-protein interactions at timescales of nanoseconds to seconds. These include Förster resonance energy transfer (FRET),^16–20^ hydrogen/deuterium exchange mass spectrometry (HDX-MS),^21–25^ electron paramagnetic resonance (EPR) spectroscopy,^26–30^ and nuclear magnetic resonance (NMR) spectroscopy.^31–34^ Alongside static structures of GPCRs in unbound, ligand-bound, and transducer-bound states, these experiments have revealed the helix movements that occur upon GPCR activation, specific structural features that mediate these movements and the possibility of intermediates during the transition between active and inactive conformations.

Over the past decade, advancements in crystallization techniques and the design of stable GPCR fusion protein constructs have allowed for the structures of several GPCRs to be solved.^35^ Structures of the GPCRs, however, do not provide information on ensemble features. However, they can serve as an excellent starting point to be used in conjunction with structure-based methods capable of investigating conformational flexibility. In this work, we employ an Ising-like statistical mechanical model termed the Wako-Saitô-Muñoz-Eaton (WSME) model,^36,37^ which has been very successful in capturing the folding mechanisms and conformational landscapes of water-soluble proteins, to explore the structural-thermodynamic hallmarks underlying the GPCR architecture in the ligand-free form. Despite the simplicity of the approach, we not only predict many known features of GPCRs but also provide a detailed view of their complex conformational landscapes for the first time, which can be used in conjunction with experiments to explore native ensemble heterogeneity, populated substates and intermediates, activation mechanisms and allostery.

## Methods

### GPCR Database

Sixty-seven high-resolution structures were downloaded from the GPCR-EXP database^38^ (experimentally solved GPCR structures) out of which only those structures that consisted primarily of the transmembrane domains were selected. In other words, those structures with large intra- or extra-cellular domains were discarded as the hydrophilic environments in which such domains exist and the hydrophobic environment within the lipid bilayer cannot be modeled simultaneously using a single dielectric constant (*vide infra*). Structures with large intra- or extra-cellular loops that are not amenable to modeling using the Robetta server^39^ were also discarded. The pruning eventually resulted in a database of 45 GPCRS - 41 from humans and one each from bovine, mouse, rat, and viral taxa (Supporting Information Table S1). Any missing residues or short loops were again modeled using the Robetta server. The sequences of these missing segments, including the third intracellular loop (ICL3), which is often replaced with a fusion protein to facilitate crystallization, were obtained from UniProtKB. Missing N- and C-terminal segments in structures obtained from truncated GPCR constructs were not modeled. Sequence modifications already present in the original PDB structures, including thermostabilizing mutations, were left unaltered. Of the mammalian GPCRs, 39 belong to class A, the rhodopsin-like receptor family. Classes B, C, and F are represented by the calcitonin receptor, two metabotropic glutamate receptors, and Frizzled-4, respectively. These 45 structures are of the GPCRs in their inactive states, bound only to a ligand (agonist or antagonist) on the extracellular side. Note that we use the term ‘inactive’ to refer to ‘non-G-protein-bound’ structures. Rhodopsin is considered only in the retinal-free form. The database additionally contains the structures of 8 GPCRs in their active conformations, with their transmembrane helices having undergone rearrangements characteristic of the active states through the binding of both an agonist on the extracellular side and a transducer protein or an antibody on the intracellular side.

### Wako-Saitô-Muñoz-Eaton (WSME) Model

The WSME model is a structure-based statistical mechanical model that employs a Gō-like treatment for its energetics, *i.e*. only those contacts or interactions that are present in the native structure are assumed to influence the folding mechanism, and is therefore entirely native-centric in its description (non-native interactions are not considered). While the model is explained in detail in many works before,^40–42^ it is briefly discussed here. In the classic WSME treatment,^36,37^ every residue is assumed to sample two conformational substates, folded (represented by the binary variable *1*) and unfolded (*0*), resulting in *2^N^* microstates or conformations for an *N*-residue protein. In the current treatment, we employ a computationally less-intensive approach wherein only stretches of consecutive residues, termed blocks, are considered as the folding unit.^40^ For example, for a 300-residue protein assuming a block-size of 3 will reduce the number of folding units to 100 blocks, instead of 300 units. Furthermore, the instantaneous ensemble is considered to be constituted from single stretches of folded blocks (single sequence approximation or SSA), two stretches of folded blocks (double sequence approximation or DSA), and DSA allowing for interactions between the folded islands (DSA with loop, or DSAw/L).^40,43^ One can imagine each of these microstates to be an array of strings with *1*s and *0*s defining the regions that are structured and hence the extent of structure; the constraint is that there can be at most only two islands of ones (DSA or DSAw/L) (Figure S1). The latter DSAw/L approximation is critical as it allows for two folded islands to interact with each other if they do so in the folded structure – the precise stability of such microstates will be determined by the relative balance between stabilizing interactions within and between the folded islands and the entropic cost of fixing residues in the intervening disordered loop. The resulting bWSME model (b standing for block) has been tested on multiple proteins and has provided insights into folding mechanisms and function in an experimentally consistent manner.^40–42^ In the current work, a fixed block length of 4 (i.e. four consecutive residues and ensuring that the block definition does not span two different secondary structure elements) has been employed for all GPCRs. The total number of microstates within the model approximation is the sum of the binomial coefficients 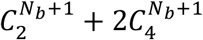 where *N_b_* is the total number of blocks (Figure S1).

The total partition function (*Z*) for the bWSME model is calculated as:

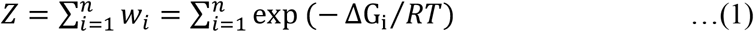

where *n* is the total number of microstates (i.e. microstates defined by SSA, DSA and DSAw/L), *w_i_* is the statistical weight of state *i, R* is the gas constant and *T* is the temperature. The free energy of every microstate with structure between and involving blocks *p* and *q* (*p, q*) is

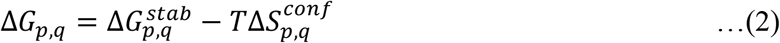

The stabilization free energy 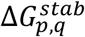 includes contributions from van der Waals interactions (interaction energy *ξ* for the vdW contacts identified from the ligand-free native structure with a 5 Å cut-off including nearest neighbors), charge-charge interactions at pH 7.0 without a distance cut-off via the Debye-Hückel formalism, and a contacts-scaled implicit heat capacity term (Δ*G_solv_*, calculated as the heat capacity change per native contact 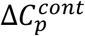 that is fixed to −0.36 J mol^−1^ K^−1^ per native contact).^41,44^ Charge-charge interactions are calculated with an effective dielectric constant of 4 to account for the low dielectric environment of the membrane environment,^41,45,46^ similar to the approach employed for studying water-soluble proteins where the solvent is considered as a uniform high dielectric continuum. The unstructured loops connecting the transmembrane helices are partially exposed to the solvent; we account for the disordered nature of the loops by attributing a higher entropic penalty for ordering (see below) thus recapitulating expectations from the structure. The role of disulphide bridges were not explicitly considered and the GPCRs were assumed to be in the reduced form to be amenable for characterization by the bWSME model.

The entropic penalty incurred for fixing all blocks in the folded conformation for the microstate (*p, q*) is given as,

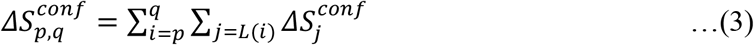

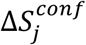 is the entropic penalty for fixing the residue *j* in the folded conformation (fixed at −10 J mol^−1^ K^−1^ per residue) while *L(i)* includes the set of residues within block *i*. An excess entropic penalty (ΔΔS) of −6.1 J mol^−1^ K^−1^ per residue is additionally assigned to residues identified as coil by STRIDE^47^ (mostly the loop regions connecting TM helices).^48^ The entropic penalty of fixing a proline residue in the native conformation is considered to be 0 J mol^−1^ K^−1^, owing to its limited conformational flexibility. Partial partition functions are calculated by grouping microstates with a specific number of structured blocks from which the one-dimensional free energy profiles are generated. For example, the effective free energy of states with 30 structured blocks is calculated from:

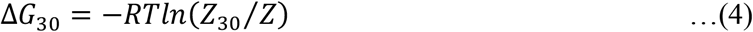

where *Z*_30_ is sum of statistical weights of states with 30 structured blocks. For comparison between proteins of different lengths (*N*), the fraction of structured blocks calculated by normalizing the number of structured blocks by the maximum number of structured blocks (*N_b_*) in the protein, was employed. A similar calculation was employed to construct 2D landscapes for specific combination of blocks structured in the N- and C-terminal halves of the protein. The folding probability of a specific block/residue *i* is calculated from:

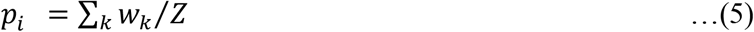

where k runs over all microstates in which residue *i* is folded. From equation (5), the stability (*s*) of residue *i* in the context of the structure is:

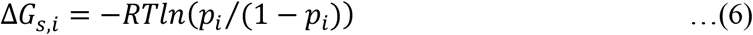

and the mean stability of a secondary structure (*ss*) element as:

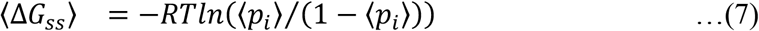

where the square brackets average over all residues corresponding to a specific secondary structure element. The heat capacity profiles were estimated *via* the derivative expression:

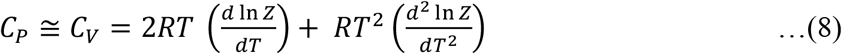

For the database of GPCRs in Table S1, only the van der Waals interaction energy (*ξ*) was adjusted such that the resulting heat capacity curve has a peak heat capacity at 333 K (the melting temperature, *T_m_*). In specific cases, where two heat capacity peaks were present, the *ξ* was adjusted such that the trough between the two peaks falls at 333 K. For GPCRs that have had structures in both active and inactive states experimentally determined, different *ξ* values were employed such that the free-energy differences between the folded and unfolded states of a receptor in both its active and inactive states were equal (*i.e*. under conditions of iso-stability). Note that ligands are not considered in the analysis and only the information from the polypeptide is employed to generate an ensemble of states and their relative statistical weights. The PDB structures employed, the model outputs, including the scripts for analyzing them are available at: https://github.com/AthiNaganathan/GPCR-Landscapes.

## Results

### Sequence and Structural Diversity in the GPCR Database

Sequences corresponding to each of the 45 GPCR structures were used to generate a multiple sequence alignment (MSA) using ClustalW.^49^ A percentage sequence similarity matrix was computed from pairwise similarities between the sequences in the MSA. Most GPCR sequences in our database exhibit low pairwise similarities, yielding a mean similarity of 9.8% (σ = 2.4%) between non-identical sequences. A high sequence similarity (59.2%) is observed between the two metabotropic glutamate receptors (GPCRs 11 and 35). Similarly, several other GPCRs belonging to the same receptor subfamily display higher than average pairwise sequence similarities. These include the chemokine receptors (GPCRs 2, 4, 20, 24, 36, and 44), the β-adrenoceptors (GPCRs 3 and 7), the proteinase-activated receptors (GPCRs 6 and 21), the opioid receptors (GPCRs 8, 9, 10, and 19), and the melatonin receptors (GPCRs 41 and 42). The structural diversity of the GPCR dataset was also probed by computing pairwise root-mean-square deviations (RMSD) between the structures using the Dali protein structure comparison server.^50^ Although the receptors show a high level of sequence divergence, structural similarity is found to be quite high with a mean pairwise C_*α*_-RMSD of 3.1 Å (σ = 0.6 Å). The only standout GPCR being the β_2_-adrenergic receptor (β_2_AR; GPCR7) that displays high pairwise RMSD values against all other structures used, including the β_1_-adrenoceptor with which it shares a subfamily (RMSD 4.8 ± 0.6 Å). The sequence-structure analysis effectively reveals that the dataset chosen is diverse enough to explore generic trends in the folding-conformational behaviors of GPCRs.

### Folding Free-Energy Profiles and Intermediates

The bWSME model was used to iteratively generate heat capacity curves at different values of the vdW interaction energy per native contact (*ξ*) while keeping every other parameter constant (Figure S2 and Table S1). The magnitude of ξ that resulted in an apparent melting temperature (*T_m_*) of 333 K was selected (see Methods). This was done to ensure that the energy scales match melting temperatures of mesophilic proteins, which is ~333 K. Note that the melting temperature of GPCRs are generally lower than 333 K and are expected to be different depending on the GPCR identity. The higher *T_m_* value assumed here is to ensure that the predictions constitute the lower limit of conformational heterogeneity. The effective mean of *ξ* across the 45 proteins is −48.9 ± 2.76 J mol^−1^ per native contact, indicating that none of the structures exhibit unique differences in packing that could contribute to extreme *ξ* values. In fact, the magnitude of *ξ* matches that of the 6-12 Lennard-Jones interaction potential between two carbon atoms (−46.1 J mol^−1^ at 6 Å) calculated from atomic-level force-field parameters.^51^

One-dimensional free-energy profiles (1D FEPs) were then generated at 333 K as a function of the fraction of structured blocks, which is a natural coordinate for the WSME model (Methods). The complexity of the profiles are better observed at 333 K as the favorable gradient towards the folded state at say, 298 or 310 K, obscures the features. The high sequence diversity observed in our dataset is expected to contribute to large differences among the free energy profiles and this is indeed the case. For example, some GPCRs present two-state-like free energy profiles with a large thermodynamic barrier between the folded and unfolded states (Figure 1B). These include Free fatty acid receptor 1 (GPCR22, Supporting Table S1) and C-C chemokine receptor type 5 (GPCR24). Others, like P2Y purinoceptor 1 (GPCR14), Orexin receptor type 1 (GPCR18) and Prostaglandin E2 receptor EP3 subtype (GPCR40), exhibit multi-state profiles containing numerous intermediates. The free energy profile of Free fatty acid receptor 1 (GPCR22) features a large free energy barrier between the folded and unfolded minima and a narrow folded-state minimum. On the other hand, the free energy profile of Adenosine receptor A1 (GPCR23) features a broad folded-state minimum. Type-2 angiotensin II receptor (GPCR25), Substance-P receptor (GPCR37) and Calcitonin receptor (GPCR43) display free energy profiles that are largely flat, suggestive of a loosely coupled structural scaffold. The positions of major intermediates on the free energy profiles also differ between different GPCRs. While the free energy profile of Sphingosine 1-phosphate receptor 1 (GPCR5) features intermediates that precede the major folding barrier, *β*_1_ adrenergic receptor (GPCR7) populates intermediates after the major folding barrier. In many cases, the native ensemble is not defined by a single state, but by a collection of sub-states either over a barrier or as a continuum of states, and this can be seen in Rhodopsin, P2Y purinoceptor, Neurotensin receptor type 1, Cannabinoid receptor 2 and Thromboxane A2 receptor.

We estimate the number of intermediates *via* a simple heuristic: a local minimum on the 1D FEP is considered to be a partially-structured intermediate if it was separated from its neighboring minima by free energy barriers of at least 1 *RT*. According to this criterion, the 1D FEPs of most GPCRs in the dataset are found to contain at least 2-3 intermediates (Figure 1C and Table S2). P2Y purinoceptor 1 (GPCR14) and Orexin receptor type 1 (GPCR18) populate the highest number of intermediates (6). It is important to note that the observed heterogeneity in the 1D FEP is only a lower limit. This is because the reaction coordinate, the number of structured blocks, lumps together millions of microstates to construct partial partition functions and hence folding free energy profiles.

Though the reliability of the WSME model predictions have been validated in numerous water-soluble proteins, it is not clear if they equally applicable to membrane associated systems. The robustness of the bWSME model energy function is showcased by studying two bacterial membrane proteins, GlpG and PagP (Table S3), whose folding mechanisms have been re-constructed from experiments. GlpG, which consists of six transmembrane α-helices, folds primarily through a mechanism that involves the folding of the entire N-terminal region of the protein before the C-terminal region folds (Figure S3A-S3D). This is in close agreement with data from single-molecule unfolding experiments and mutational analysis, which indicate that the C-terminal region is more unstable and the presence of a N-terminal biased folding nucleus.^52^ The β-barrel membrane protein PagP folds via an intermediate in which parts of both the N- and C-terminal regions are structured, with a higher probability of C-terminal structure.^53^ This observation is again in good agreement with our model, which yields a two-dimensional free energy surface with a significant local minimum in which both the N- and C-terminal regions are partially structured with a higher structural disposition towards the C-terminal strands (Figure S3E-S3H). The agreement of the model predictions with experimentally-constructed folding mechanisms thus attests to the robustness of our method and the uniform dielectric constant employed for studying membrane proteins. We delve into the thermodynamic architecture of GPCRs in the sections below.

### Helix Stabilities and Conformational Plasticity

To examine how stability determinants are distributed across the GPCR structures, the folding probability of every residue in the protein was calculated at 310 K by summing up the statistical weights of microstates with a specific residue folded and their relative contribution to the total partition function (Methods and equations 5–7). These residue folding probabilities were then used to compute the average stability of residues within each of the seven transmembrane helices for all 45 GPCRs (Figure 2A). Note that this calculation allows for the estimation of the helix stability in the context of the structure considered and not in isolation. We find that TM3 is the most stable of all the helices, with TM1 being the least stable. Thus, without explicitly considering the disluphide bridge between the first and second extracellular loops (ECL1 and ECL2), the model is still able to predict the larger stability of TM3. The stabilities of TM1, 6 and 7 vary substantially in the dataset studied and, in some GPCRs, these helices are unstructured even in the native ensemble (positive helix stability values in Figure 2A).

**Figure 2.**
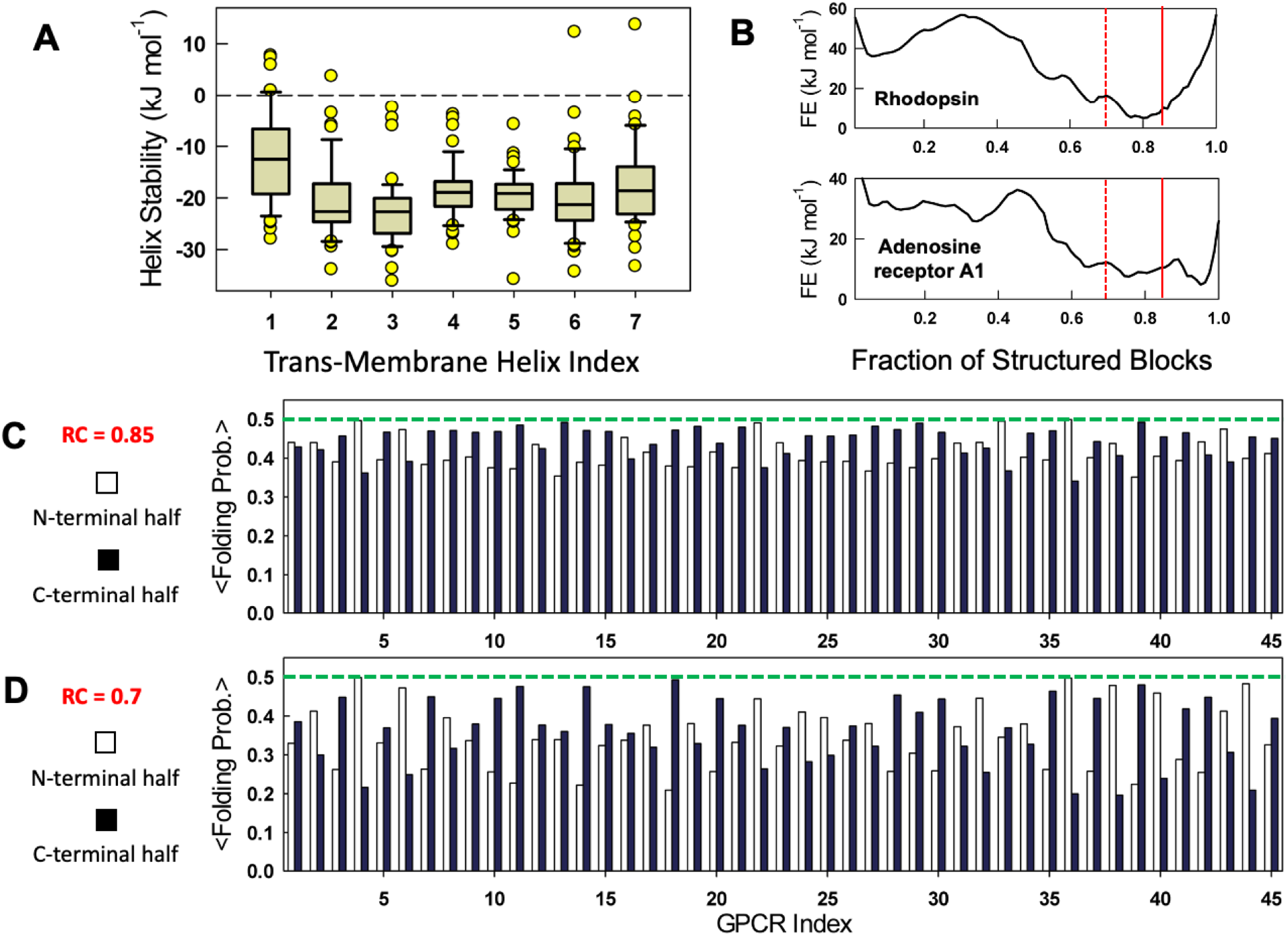
Helix stabilities and conformational plasticity at 310 K. (A) Box plot of individual TM helix stabilities. (B) One-dimensional free energy profiles of two representative GPCRs. The vertical dashed lines signal the reaction coordinate (RC) values of 0.85 (continuous red) and 0.7 (dashed red), respectively. (C, D) Protein regions that unfold first as one moves from the right to the left on the free energy profiles are illustrated by dividing the GPCRs into the N-terminal half (TM helices 1-3) and C-terminal half (TM helices 4-7). Panels C and D plot the means folding probability of the N- and C-terminal halves of the structure at the indicated reaction coordinate values.

As a second step, one-dimensional free energy profiles were constructed at 310 K as a function of the reaction coordinate, the number of structured blocks (Figure 2B). Regions of the GPCR that unfold first or are partially structured in the native ensemble are identified by choosing two specific regions on the reaction coordinate (RC=0.85 and RC=0.7; vertical lines in Figure 2B) and plotting the probability of structure in the N-terminal half (〈*p_f,N_*〉) versus C-terminal half of the structure (〈*p_f,C_*〉) (Figure 2C, 2D). The former accounts for the first three TM helices while the latter accounts for the TM helices 4-7. At RC=0.85 which corresponds to the near-fully folded native ensemble, both the N- and C-terminal halves are already partially structured in the majority of GPCRs with 〈*p_f,N_*〉 of 0.41 compared to 〈*p_f,C_*〉 of 0.44 on average (Figure 2C). Importantly, 30 of the 45 GPCRs exhibit more unfolding in the N-terminal half compared to the C-terminal half. Minor perturbations can be mimicked by observing the stability patterns at RC = 0.7 where the protein is marginally more destabilized (Figure 2D). Under these conditions, both the protein halves are similarly unfolded on average across all proteins (〈*p_f,N_*〉 of 0.34, and 〈*p_f,C_*〉 of 0.36), with the distribution flipped in favor of more unstructured C-terminal halves in 27 GPCRs (Figure 2D). Given that TM1, 6 and 7 exhibit lower stabilities compared to other helices, it is likely that these regions sample partially structured states in the native ensemble. Topologically, this conformational behavior is expected as TM1 is the most weakly packed of all helices, interacting only with TM2 in most proteins and also with TM7 in some (Figure 1A). On the other hand, TM7 is relatively more packed, directly interacting with all helices except TMs 4 and 5, and hence it is less likely to sample unstructured states compared to TM1.

To investigate this further, 2D free energy landscapes were generated for all GPCRs at 310 K, with the number of structured blocks at the N- or C-terminal region as coordinates. Such a 2D landscape has been particularly successful in capturing functionally relevant substates in multiple large water-soluble proteins.^42,54,55^ For example, consider the free-energy landscapes of Rhodopsin (GPCR1), the *β*_2_AR (GPCR3) and the Kappa-type opioid receptor (GPCR8) at 310 K (Figure 3). The native ensembles are broad in the GPCRs considered, but with differences in the extent and nature of structure. The Rhodopsin free energy surface (Figure 3A) is indicative of a continuum of states in the native basin (states *a* and *b*), while the states *b* and *c* in the *β*_2_ AR (Figure 3B) and all labeled states in the Kappa-type opioid receptor (Figure 3C) are intermediate-like, and are populated over a marginal thermodynamic barrier. The partially structured states *c* and *a* in Rhodopsin and the *β*_2_AR, respectively, will however not have a large residence time as they appear as ‘excited states’ along the coordinate (they do not constitute a minima on the landscape). In these three GPCRs, it is TM1 that exhibits the largest degree of unfolding.

**Figure 3.**
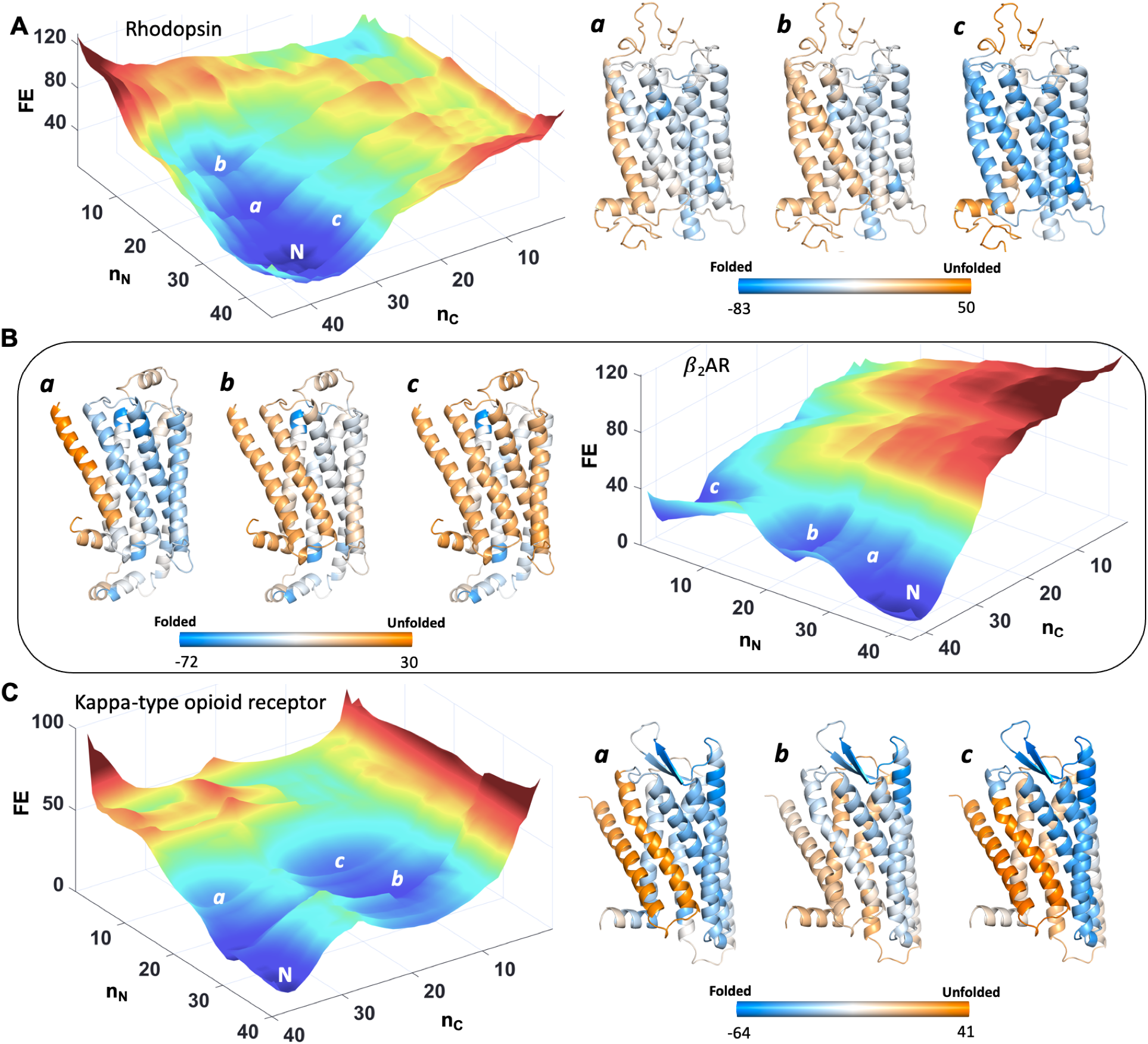
Free energy landscapes of Rhodopsin (panel A), *β*_2_AR (panel B) and Kappa-type opioid receptor (panel C). The free energy values and the color bars are in units of kJ mol^−1^. The structures are color coded according to the color bars with light blue and dark orange representing fully folded and fully unfolded conformational status. Intermediate free energy values colored in white represent partial structure. The major conformational states *a, b*, and *c* are shown adjacent to the free energy landscapes with N representing the folded native ensemble.

In Rhodopsin state *a* (Figure 3A), one would expect the unfolding of TM1 to not affect the adjacent helices, but it is clear that the free energy of folding (equation 6) of almost all the helices are perturbed – they should be in the dark blue color range (more folded) but instead fall in the region between cyan and white (partially unfolded). This is a consequence of the fact that a loss of interactions between TM1 and TMs 2 and 7 in turn destabilizes the TMs adjacent to them but to a lesser extent, similar to the effect of mutations on protein structure.^56^ State *c* in Rhodopsin is characterized by fully folded TMs 1-4 while TMs 5-7 are partially structured. In the *β*_2_AR, unstructured TMs 1-2 are the predominant substates (states *a* and *b* in Figure 3B), similar to the state *a* in the kappa-type opioid receptor (Figure 3C). Additionally, the substate *a* in *β*_2_ adrenergic receptor exhibits partial structure in TMs 6 and 7 (white in Figure 3B), which mirrors experimental observation of substates involving significant mobility in the same set of helices.^57^ Furthermore, partial structure in TMs 1, 2, 6 and 7 of the kappa-type opioid receptor promote the population of an intermediate *c* that has only TMs 3, 4 and 5 folded (Figure 3C). To summarize, it appears that while partial unfolding of TM1 is a dominant substate in GPCRs, there can be substantial variations in the nature of the states populated and their relative populations.

### Anisotropic Distribution of Coupling Free Energy Magnitudes

GPCR activation mechanisms are dependent not just on the thermodynamic stabilities of individual helices (in the structural context) but also the extent to which these stabilities are modulated via altered structural patterns and contacts *between* helices on ligand binding. A precise understanding of this could be gleaned by computing the extent to which the different regions of the protein are thermodynamically coupled to each other.^58,59^ We calculate coupling free energies between residues from the bWSME model by grouping the ensemble of microstates into four different sub-ensembles for every residue *i* (Figure 4A): ∑*p_i_f_j_f__* sums over the probabilities of all states in which both residues *i* and *j* are folded, ∑*p_i_f_j_u__* sums over probabilities of states in which residue *i* is folded and *j* is unfolded, and similarly for ∑*p_i_u_j_f__* and ∑*p_i_u_j_u__*.^41^ From these groupings, one could calculate positive (Δ*G_+_*), negative (Δ*G_−_*) and effective (Δ*G_C_*) coupling free energies^41,60^ between different residues using:

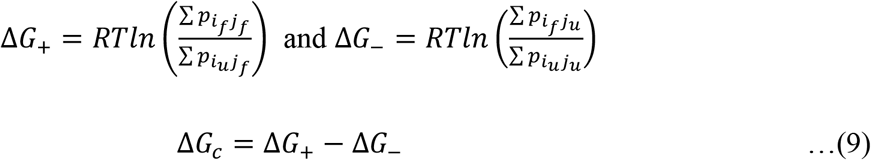

**Figure 4.**
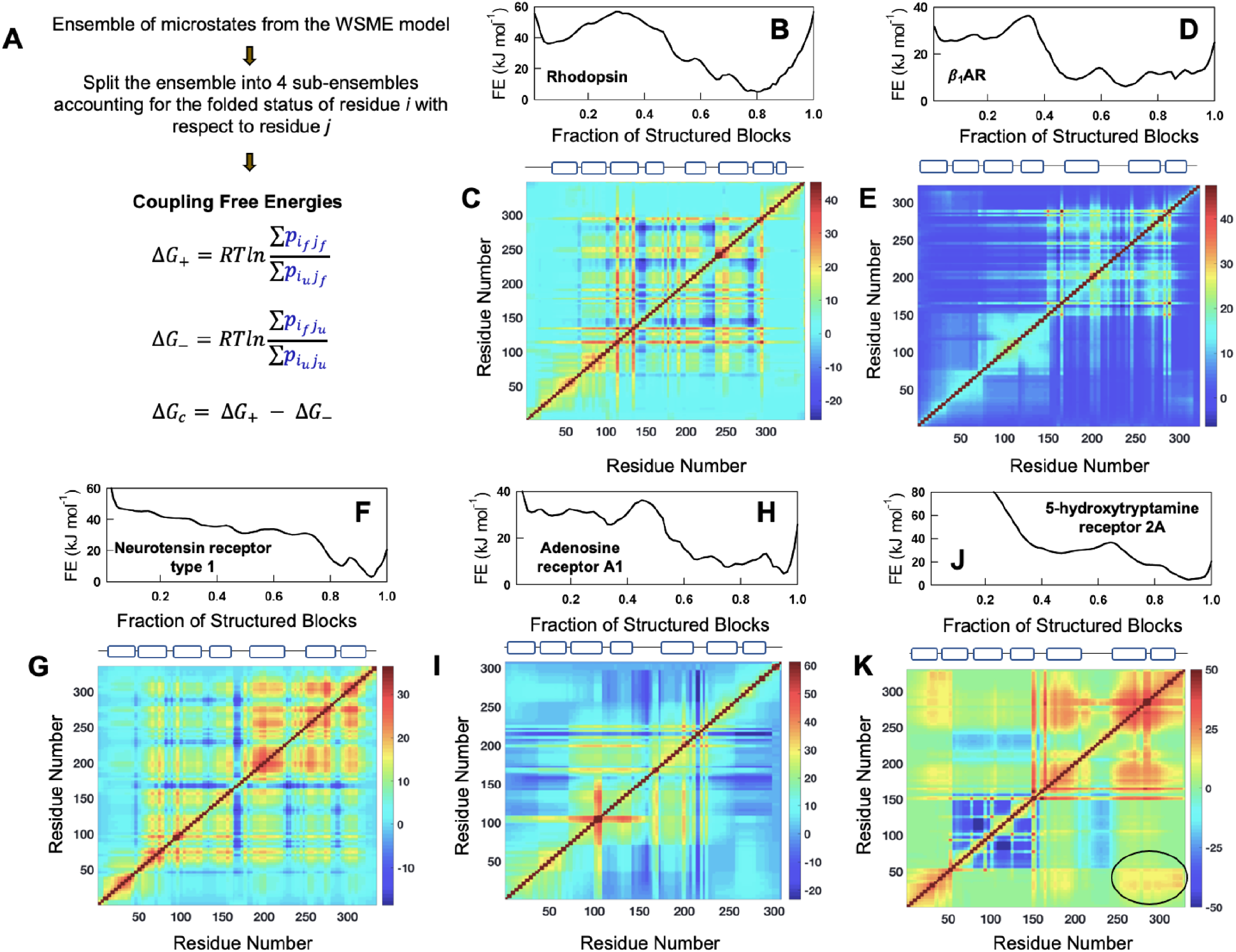
Thermodynamic architecture of GPCRs. (A) A flow-chart describing the methodology employed to calculate coupling free energies from the microstate populations. (B, D, F, H, J) Onedimensional free energy profiles at 310 K. (C, E, G, I, K) The corresponding effective coupling free energy (Δ*G_C_*) matrices. The color bars are in the spectral scale and in units of kJ mol^−1^, with dark red and dark blue representing strong and weakly coupled residues, respectively. The cartoons on top indicate the position of the secondary structure elements along the sequence. The oval in panel K highlights that the TM helices 1 and 7 are strongly coupled in 5-hydroxytryptamine receptor 2A.

Positive coupling free energies quantify the extent to which residues *i* and *j* are coupled via direct interactions or through long-range interactions in the native ensemble while the negative coupling free energies quantify the extent to which lack of spatial proximity, unfavorable interactions or large conformational entropy decouples specific structural regions from others. The balance between the two terms results in effective coupling free energies – residues that present lower effective coupling free energies are typically located in functional or dynamic regions of the structure as shown for multiple proteins in a recent work.^41^ Importantly, coupling free energies can be calculated for every residue with respect to every other residue (and hence a square matrix can be constructed), revealing insights into the distribution of stabilization free energies in the structure.

Given the range of GPCR free energy profiles and individual helix stabilities, the coupling maps are expectedly not uniform across the GPCR dataset. For instance, in Rhodopsin (GPCR1), the broad native well Figure 4B is a manifestation of minimal coupling between N- and C-terminal regions (note the cyan scale in Figure 4C). In the *β*_1_AR (GPCR7), the native ensemble is not composed a single state but a continuum of conformations (Figure 4D) which results from weak inter-residue coupling between the majority of residues in the protein (sea of blue in Figure 4E). More complex patterns are also evident for Neurotensin receptor type 1 (Figure 4F, 4G) and Adenosine receptor A1 (Figure 4H, 4I), with distinct coupling free energy patterns. 5-hydroxytryptamine receptor 2A (GPCR31), meanwhile, exhibits strong coupling between residues in its C-terminal region with particularly strong inter-helical coupling between TM1 and TM7 not seen in the other members discussed here (circled regions in Figure 4K).

The diverse patterns in Figure 4 are better observed by mapping the residue-averaged coupling free energies (〈Δ*G_C_*〉, i.e. averaging along the dimensions of the symmetric matrices in Figure 4) onto the three-dimensional structure. It can be seen that the coupled residues (different shades of magenta) are not uniformly distributed throughout the structure but are localized to specific regions in the protein (Figure 5A). The 〈Δ*G_C_*〉 values were Z-scored to account for intrinsic differences in the range of coupling free energies, and residues that exhibit a Z-score greater than one were labelled as strongly-coupled. The fraction of strongly coupled residues (f_c_) thus calculated vary for the GPCRs but are constrained within 25% (Figure 5B). In fact, an analysis of 25 water-soluble proteins (SPs) found that they exhibit a mean fc of ~16%,^41^ while the GPCR dataset exhibits a marginally lower value of ~13% (Figure 5C). Thus, despite the membrane bound nature of GPCRs, their thermodynamic architectures do not deviate significantly from those of water-soluble proteins. The anisotropic distribution patterns of strongly coupled residues are also consistent with the observations in SPs.^41^

**Figure 5.**
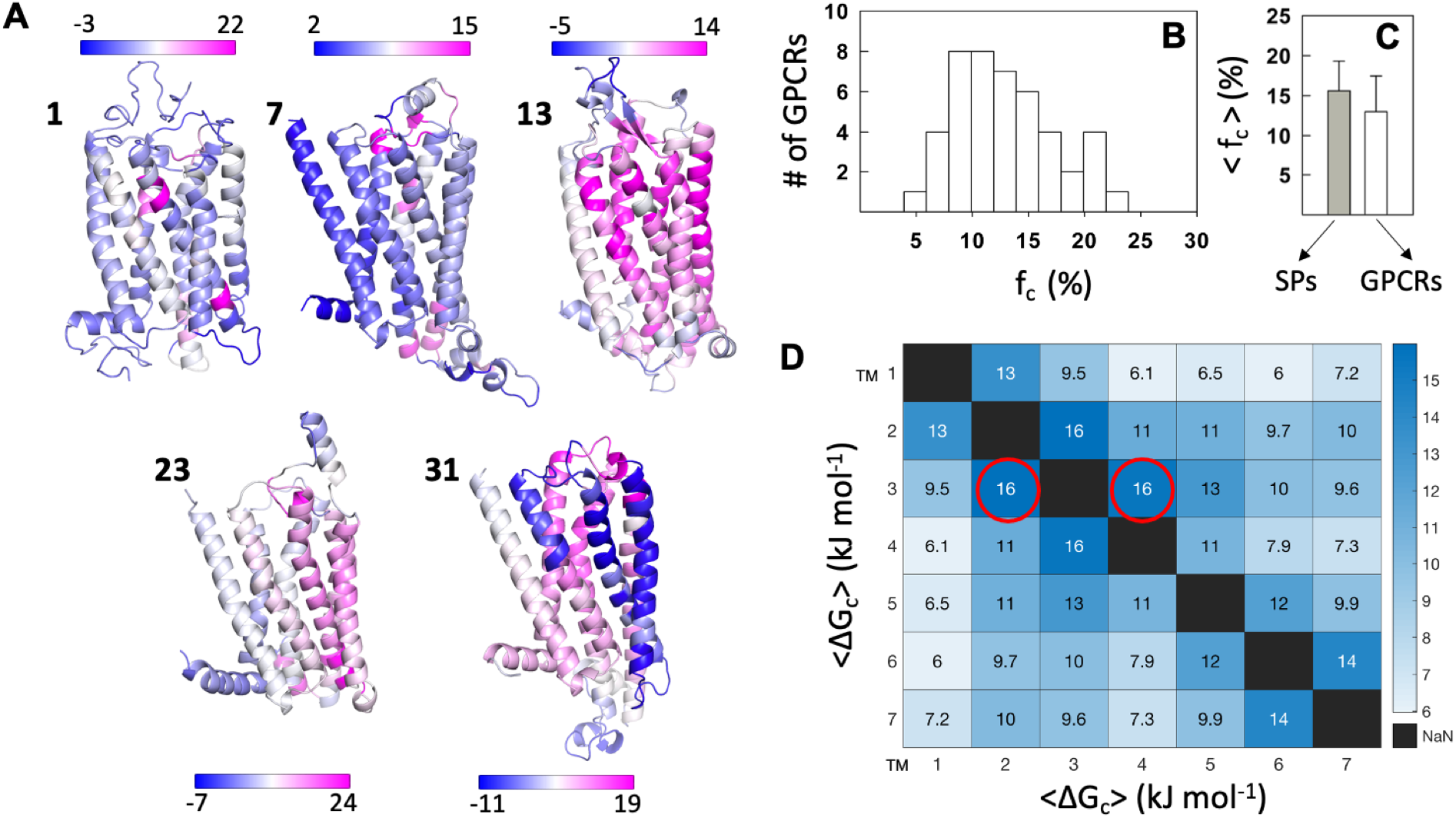
Anisotropic distribution of coupling free energies. (A) The mean effective coupling free energy from the matrices in Figure 4 are mapped on the structure of GPCRs. The GPCR indices are provided for reference. Color bars are in units of kJ mol^−1^. Regions of the protein colored in dark magenta represent strongly coupled residues while those in dark blue are not coupled. (B) Distribution of the fraction of strongly coupled residues (f_c_) in GPCRs. (C) Comparison of the mean fraction of strongly coupled residues in GPCRs and soluble proteins (SPs). (D) Effective coupling free energies between transmembrane (TM) helices averaged over the 45 GPCRs. The mean standard deviations in 〈Δ*G_C_*〉 are of the order of 4 - 6 kJ mol^−1^, indicating significant variability. Despite this variability, TM3 turns out to be strongly coupled to every other TM helix.

To derive more generalized inferences on which regions of the proteins are more coupled than others, the average pairwise coupling between helices across all 45 inactive GPCR structures is calculated to construct the inter-TM coupling matrix (Figure 5D). Transmembrane helices located adjacent to one another are strongly coupled due to the nearest neighbor effects. TM1 and TM7, on the other hand, are weakly coupled to the rest of the structure (except to TM2 and TM5, respectively), as they constitute the termini of the protein. In agreement with Figure 2A, TM3 is the most strongly-coupled of the helices, consistent with TM3’s theorized role as a structural hub that maintains the GPCR scaffold.^61^ Furthermore, in the topological arrangement of helices, TM3 interacts with all the other helices except TM1, making this region in particular more stable and crucial to the stability and functioning of GPCRs. TM4 is strongly coupled to TM helices 2, 3, and 5, while being marginally coupled to the other helices. One standout message from the inter-TM coupling matrix is the fact that every TM is either weakly or strongly coupled to one another thermodynamically. Any modulation of the coupling free energies between a pair of helices, say by ligand binding, will necessarily affect the coupling free energies throughout the structure (*vide infra*).

### Are Active State Structures More Strongly Coupled?

GPCR activation is characterized by large-scale movements of transmembrane helices.^5,14,15^ In particular, activation causes TM6 to swing outward while TM5 and TM7 move in towards the helical bundle. This should affect the coupling free energies and hence the free energy profiles depending on the extent to which interactions are formed or broken between residues in TM3, TM5, TM6, and TM7. To understand this quantitatively, free energy profiles were generated for 8 GPCRs whose structures are available in both active and inactive states (Figure 6A-H and Table S4). The number of accessible states in the active state native ensemble are minimized in Rhodopsin (GPCR1), *β*_1_AR(GPCR7), Kappa-type opioid receptor (GPCR8) and Adenosine receptor A1 (GPCR23), i.e. the native ensemble is sharper with a narrow minimum, while no significant modulations are observed in *β*_2_AR (GPCR3) and Neurotensin receptor type 1 (GPCR13). These results are consistent with the idea that inactive GPCRs are capable of sampling a variety of conformations and that agonist binding stabilizes the active-like conformation.^5^ Particularly, the finding that *β*_2_AR samples similar set of conformations in the inactive and active states is in agreement with detailed NMR experiments.^62^ On the other hand, Mu-type opioid receptor (GPCR9) and Type 1 angiotensin II receptor (GPCR16) display a broader native ensemble in their active state, indicating that the connection between the active state and a narrower ensemble is not generalizable, at least from the perspective of 1D free energy profiles.

**Figure 6.**
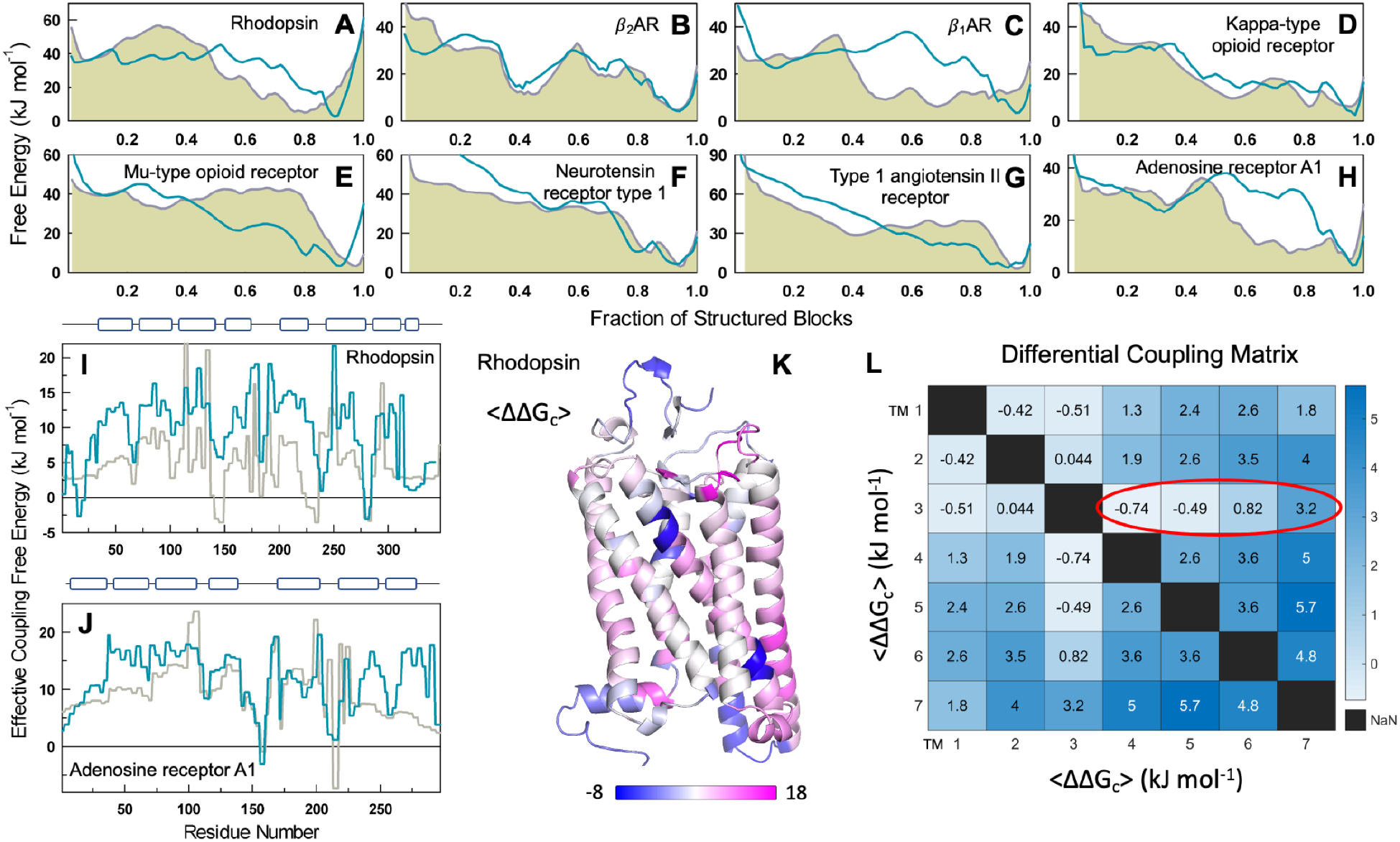
Active *versus* inactive states. (A-H) Free energy profiles derived from the inactive (gray filled) and active (dark cyan) GPCR structures at 310 K. (I, J) Effective free energies as a function of residue index for two representative GPCRs. Gray and dark cyan represent the effective coupling free energies calculated from inactive and active states, respectively. The cartoons on top indicate the position of the secondary structure elements along the sequence. (K) Differences in effective coupling free energy (active – inactive) from panel I is mapped on to the structure of Rhodopsin. (L) The differential coupling matrix (active – inactive) from the perspective of transmembrane helices. The dark red oval represents a gradation in the coupling of TM3 with the other helices on activation; this highlights the motions of other helices around TM3. The mean standard deviations in 〈ΔΔ*G_C_*〉 are of the order of 4.5 – 6.5 kJ mol^−1^, suggestive of large variations across the 8 pairs of active-inactive states.

We further computed the effective coupling free energy matrices for the active and inactive structures and averaged them along one dimension to plot them as a function of sequence index. Residues tend to be more strongly coupled to the rest of the structure in active-state structures compared to inactive-state structures on average (Figure 6I, J and Figure S4). The stronger coupling in the active form is more evident in the case of Rhodopsin (Figure 6I), Adenosine receptor A1, *β*_1_AR and Mu-type opioid receptor (Figure S4) wherein nearly all helices are stabilized. *β*_2_AR, on the other hand, displays little change in the degree of coupling though differences can be observed between and including TMs 5 and 6. The inactive structure is, however, more coupled in the Neurotensin receptor type 1 and Type 1 Angiotensin II receptor. The differential wiring of the interaction network in each of the protein potentially contributes to the differences we observe from the perspective of the coupling free energies, highlighting the intrinsic malleability of the contact-network in GPCRs.

Finally, the difference between the effective pairwise coupling between transmembrane helices in active and inactive structures was computed for the 8 GPCRs to generate the differential coupling matrix (Δ*G_c,active_* – Δ*G_c,inactive_*; Figure 6L). The mean pairwise thermodynamic coupling between most helices increases upon GPCR activation, despite the averaging across 8 GPCRs. The standard deviations are quite large, however, indicating that no two activated structures contribute to similar changes in the coupling free energies. Despite this, TM3, the most stable and most strongly-coupled of the helices in the inactive state, exhibits a gradation in the coupling differences between active and inactive state structures. While it is more strongly coupled to distant helices (TM6 and TM7) in the active state, coupling with TM1, TM4, and TM5 decreases upon GPCR activation.

### Alanine-Scanning Reveals Long-Distance Thermodynamic Connectivity in Rhodopsin

The comparison of active-inactive structures shows that the perturbations induced by activation can be pervasive and modulate long-range structural features. The extent to which the binding of a ligand influences a distant site could be potentially studied for every GPCR in the presence of agonists and antagonists. However, the WSME model does not include the atomic detail necessary for detailed modeling of subtle effects at the level of chemical interactions between the protein and the ligand, a feature that likely determines the differential effect of ligands on GPCR structures. Moreover, the model cannot be employed to reproduce the experimental protein-ligand dissociation constant (as of now) which is a necessary first step towards understanding ligand binding effects.

An alternative is to explore the extent to which every residue is coupled to every other residue in the folded conformational ensemble by performing alanine-scanning mutagenesis. Alanine substitutions at different positions will affect the interaction network to different levels depending on the immediate environment of the mutated residue, and in comparison with the WT connectivity matrix, one can extract the extent of coupling to a distant site. We consider only the positive coupling free energy (Δ*G*_+_) for this calculation, as it carries information on the states that harbor coupled residues in the native ensemble, while not considering the decoupled residues or microstates in the unfolded ensemble. For every mutant, a Δ*G*_+,*Mut*_ matrix (*N* × *N* matrix where *N* is the number of residues in the protein) is generated and referenced to the WT matrix (Δ*G*_+,*WT*_) to arrive at the differential positive coupling matrix (ΔΔ*G*_+_) (Figure S5), which is averaged across all pairwise sites to generate the vector 〈ΔΔ**G*_+_*〉 (dimension *N* × 1) (Figure 7A). The latter carries information on the extent to which every residue is perturbed including the direction of perturbation – positive and negative change are indicative of stronger and weaker coupling, respectively – for a given alanine mutation. If the alanine mutation is performed across *m* sites on the protein, the resulting 〈ΔΔ**G*_+_*〉 matrix (dimension *N* × *m*) is employed to generate the mean *μ* and standard deviation *σ* of the mutational response (MR, dimension *N* × 1).

**Figure 7.**
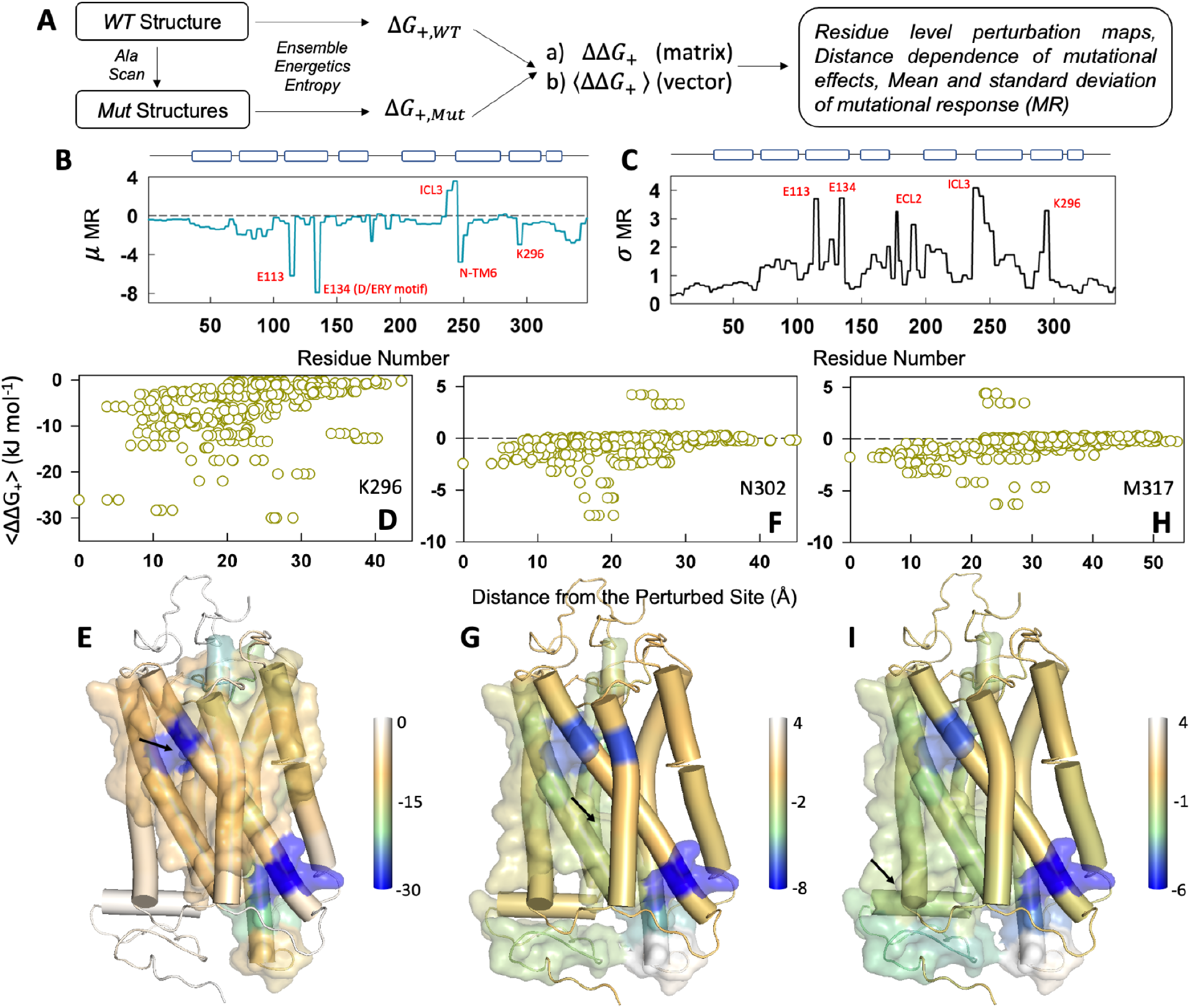
Alanine-scanning reveals long-range communication in Rhodopsin. (A) Protocol employed for *in silico* alanine-scanning mutagenesis studies from the perspective of the WSME model. (B, C) Mean (panel B) and standard deviation (panel C) of the mutational response (MR in kJ mol^−1^). Specific residues and structural regions are highlighted for discussion in the main text. E113 and K296 are in the retinal binding pocket while E134 forms a part of the conserved D(E)RY motif that interacts with G-protein. N-TM6 represents the N-terminal region of TM6. (D, E) Perturbation response of K296 quantified by 〈ΔΔ*G_+_*〉 as a function of C_*α*_-distance from the mutated site (panel D). 〈ΔΔ**G*_+_*〉 values are mapped on to the structure in panel E following the color code shown to the right. Residues that exhibit a 〈ΔΔ**G*_+_*〉 value stronger than the average are represented in the surface mode. The arrow points to the location of the residue that is perturbed. (F, G, H, I) Same as panels D and E for N302 (panels F, G) and M317 (panels H, I), respectively.

We carried out a large-scale in silico alanine scanning mutagenesis of Rhodopsin involving 276 sites on the protein excluding the positions containing alanine, glycine and proline. The process involved introduction of mutations using PyMol, construction of ensembles with parameters identical to that of the WT, and, following this, generation of 〈ΔΔ**G*_+_*〉 matrices (dimension 348 × 276; Figure S6). The resulting mean mutational response highlights specific protein regions whose coupling magnitudes change the most on perturbations (Figure 7B). First, the retinal-binding site residues E113 (which is a part of **E**GFF sequence block) and K296 (part of the FFA**K** block), both of which line the orthosteric site, stand out as positions that exhibit large changes on mutational perturbations. Second, the G-protein binding pocket involving the D(**E**)RY motif (E113) and the intracellular loop 3 (ICL3)/N-terminal region of TM6 exhibits high sensitivity to mutations across the structure. The same regions additionally exhibit larger standard deviations in the mutational response, indicating that the Rhodopsin structure exhibits intrinsic differences in dynamics (and hence thermodynamic coupling) depending on the location of the perturbation (Figure 7C).

The residue-level 〈ΔΔ*G*_+_〉 vector that is generated for every mutation carries information on the degree to which different residues are thermodynamically coupled and hence the extent (quantified in terms of distance) to which such perturbation effects are felt. As representative examples, we discuss three different residues – K296, N302 and M317 – that play critical roles in the functioning of Rhodopsin, and GPCRs in general.^63^ Perturbation of K296 in the retinal binding pocket (*i.e*. a K296A mutation) induces strong destabilization across the structure when compared to the WT. This can be observed as negative 〈ΔΔ**G*_+_*〉 and that modulates the folding status of residues located as far as 35-40 Å from the mutated site with the major effect within 25 Å (Figure 7D, S5). Mapping these magnitudes on to the structure (Figure 7E), it is clear that any perturbation of K296 or residues around it will naturally modulate the extent of coupling at the G-protein binding site located at the intracellular side of Rhodopsin (with residues spanning TM helices 3,5,6, and 7). This can be seen from the surface representation for strongly coupled residues in Figure 7E that spans the entire length of Rhodopsin. Thus, it appears that the ligand binding pocket is strongly and thermodynamically connected to the G-protein binding pocket. Perturbation of N302 in TM7 (from the NPxxY motif) reveals that this residue is thermodynamically coupled to a majority of residues at the intracellular side (Figure 7F, 7G). Remarkably, this connection, as represented by the surface map in Figure 7G, extends all the way to the ligand-binding pocket though the magnitude of this coupling is ~4 times lower compared to K296. The residues in helix 8 have been proposed to also interact with G-protein to enable the formation of functional complexes.^64^ While perturbation of K296 reveals little effect on helix 8, we find that perturbation of M317 located in helix 8 is sensed at the ligand binding site including K296 and the surrounding residues (Figure 7H, 7I). The lack of reciprocity (K296 versus M317, for example) reveals that conformational modulations can be fine-tuned to accommodate a ligand with different distantly located protein regions providing their feedback to the ligand binding site(s), potentially determining their unbinding rates.

## Discussion

The diverse sequence features of the GPCR family are implicitly accounted for by the WSME model energy-entropy function - sequence-structure dependent conformational entropy, charge-charge interactions and packing interactions - and the observed native ensemble heterogeneity is an emergent property of these small sequence-dependent features. One of consistent observations is the presence of kinetic traps or intermediates in the free energy profiles; these intermediates are likely a manifestation of functional requirements as shown recently for several large water-soluble proteins.^42^ True to this expectation, the functionally important TM helices (TMs 1, 6, and 7) are typically only marginally coupled to the rest of the structure, exhibit low intrinsic stability in the inactive state and are partially unstructured in the native ensemble. This conformational preequilibrium, defined as the co-existence of both fully folded and partially structured substates with varying probabilities on the conformational landscape, either over a broad native ensemble or as a series of intermediates and/or excited states, is potentially one of the reasons for the difficulty in crystallizing GPCRs. However, the precise extent of pre-equilibrium is not universally conserved – no two GPCR conformational landscapes are similar – and is likely selected based on the identity of the ligand and the required magnitude of functional readout. These aspects need to be studied on a case-by-case basis with appropriate experimental calibration of the model.

The magnitude of coupling free energies, which are second-order measures (unlike residue folding probabilities which are first-order measures), provides insights into the structural and thermodynamic architecture of proteins. The degree of coupling of different structural regions in GPCRs can be dramatically different despite their high structural similarity, showcasing the exquisite structural evolution driven by functional requirements. Specifically, such diversity is a consequence of the anisotropic distribution of stability patterns across the structure, with the distribution of strongly coupled residues (most strongly coupled residues are in TMs 2 and 3) and the fraction of strongly coupled residues (<30%) mirroring observations in soluble proteins. The central role of TM3 as a structural hub emerges naturally from the structural-ensemble-based calculation, in addition to the precise magnitude of coupling between different TM helices in the inactive state. Structural analysis of GPCRs based on consensus contacts has revealed extensive insights into GPCR structures, likely activation mechanisms and regions of proteins that are involved in activation.^5,61^ We reformulate these implicitly into the WSME model and find that many active structures sample a significantly constrained conformational space. This can be explicitly observed both in the free-energy profiles and in the resulting coupling free-energy magnitudes. While it is not possible to generalize these observations to all GPCRs given the limited dataset, the mean changes in coupling free energies follow a specific pattern wherein all TM helices are more strongly coupled to the rest of the structure in the active state compared to the inactive conformation. Though TM3 remains rigid during these conformational motions, the effective coupling with adjacent helices is modulated from negative (weaker interactions) to positive (stronger interactions) in going from TM1 to TM7.

Given the large conformational flexibility in the ligand-free GPCRs, it is tempting to speculate that the observed helix mobility and partial unfolding is required for effective binding to various agonists and antagonists and for precise control of functional outcomes. In fact, simulations involving alprenolol binding to *β*_2_AR point to two primary pathways involving either the channel between ECL2 and TM4/6/7 helices or between ECL2 and TM2/7,^65^ which was also subsequently observed long time-scale MD simulations.^66^ There is an overall consistency between simulations and the WSME model predictions. Specifically, partial structure in TMs 1/2 (state *b* in Figure 3B) and TMs 1/2/3/4 (state *c*) can open up potential crevices and channels for ligands to bind. In Rhodopsin, retinal unbinding simulations point to unbinding from a cleft between TM 4/5 or TM5/6.^67^ If one were to expect unbinding to be the reverse of binding from the principle of microscopic reversibility, then this requires structural flexibility and partial unfolding of the TM helices 4/5/6 and this is observable in states *a* and *b* in Figure 3A. The large entropic stability of inactive GPCR conformations (because of their inherently flexible nature) is therefore likely compensated by enthalpic effects through the binding of ligands in the open crevices, cavities between TM helices or in the extracellular side (agonists, antagonists, or drugs), and via G-proteins on the intracellular side.

The mechanistic basis for diseases associated with GPCR dysfunction include inactive or constitutively active receptors, under-expressed receptors, and misfolded receptors, all of which arise due to mutations distributed across the structure. It is conceivable that mutations modulate the number and nature of intermediates or many of the minor excited states, thus influencing foldability and half-life. Such modulations could appear as differential coupling patterns in the communication network and hence manifest as allosteric effects, subtly determining the binding of various agonists, antagonists and partial agonists. This communication network is extracted by performing large-scale alanine-scanning mutagenesis on Rhodopsin as a representative example. The ligand-binding pocket is found to be strongly connected to the G-protein binding site, as evidenced by large differences in positive coupling free energies when a mutation is introduced at the ligand binding site (Figure 7), similar to the results of sequence-based statistical coupling analysis.^68^ Given the large-scale connectivity map, it appears that binding of ligands is precisely coordinated by not just the binding site residues and ‘microswitches’ but also the folded status of many residues far from binding pocket, a feature that likely determines the differences in affinity to agonists, antagonists and partial agonists. We would like to note that the perturbation method does not reveal the different ‘communication routes’ nor the fluxes through them, but reports on the extent to which a distant site is perturbed and the magnitude of perturbation. Importantly, the resulting changes in positive coupling free energies are a manifestation of the differences in the underlying distribution of states (as exemplified by equation 9) which when mapped on to a single structure reveal distance-dependent effects. The nature and the strength of ligand binding (which is effectively a perturbation to the binding site) could therefore determine the functional output by restricting the accessible conformational space as seen in the energy landscapes of activated receptors. A similar conformational feature has been recently demonstrated in the diverse class of large nuclear receptor ligand binding domains,^55^ indicating that ‘conformational selection’ and subsequent enthalpic compensation (via drug-protein contacts or GPCR-G-protein interactions) of entropic stability could be generic features underlying the energy landscapes of proteins and, hence, function.

Membranes, which are implicitly treated in the current approach as a low dielectric continuum, and their specific composition are known to affect the conformational features of GPCRs.^7,12^ Would lipids enhance or reduce the observed heterogeneity? Since the interactions will membrane components and cholesterol will stabilize specific parts of the structure, they will likely contribute to the population of additional states with lifetimes dependent on the strength of interaction. Naturally, this would enhance the ruggedness of the conformational landscape. This expectation has already been borne out in simulations involving water-soluble proteins with non-native interactions.^69,70^ Thus, the GPCR conformational complexity presented here is likely a lower estimate, with lipids, cholesterol, ions and pH modulations further tuning the equilibrium of states.

## Reporting summary

Further information on research design is available in the Nature Research Reporting Summary linked to this article.

## Data availability

The datasets generated during this study are available in the Github repository, https://github.com/AthiNaganathan/GPCR-Landscapes.

## Code availability

The code used for generating free energy profiles and coupling free energy matrices are available at https://github.com/AthiNaganathan/GPCR-Landscapes. Any scripts required for analysis are freely available on request by contacting the corresponding author.

## Abbreviations

MD: Molecular dynamics
WSME: Wako-Saitô-Muñoz-Eaton
NMR: nuclear magnetic resonance

## Acknowledgements

The authors are grateful for the support of the Science and Engineering Research Board (SERB; Department of Science and Technology, India) for the grant MTR/2019/000392 to A. N. N, and acknowledge financial support from the Ministry of Education, New Delhi (Sanction No. 11/9/2019-U.3(A)), and the Centre of Excellence in Biochemical Sensing and Imaging Technologies (CenBioSIm), Indian Institute of Technology Madras. The authors thank Anirudh Ranganathan for comments on the manuscript.

## Author Contributions

S. A. performed the simulations. S. A. and A. N. N. analyzed the data, prepared figures and wrote the manuscript.

## Additional information

**Supplementary information** The online version contains supplementary information at

**Correspondence** and request for data or codes should be addressed to Athi N. Naganathan.

