## Supporting Material for "Thermodynamic Architecture and Conformational Plasticity of GPCRs"

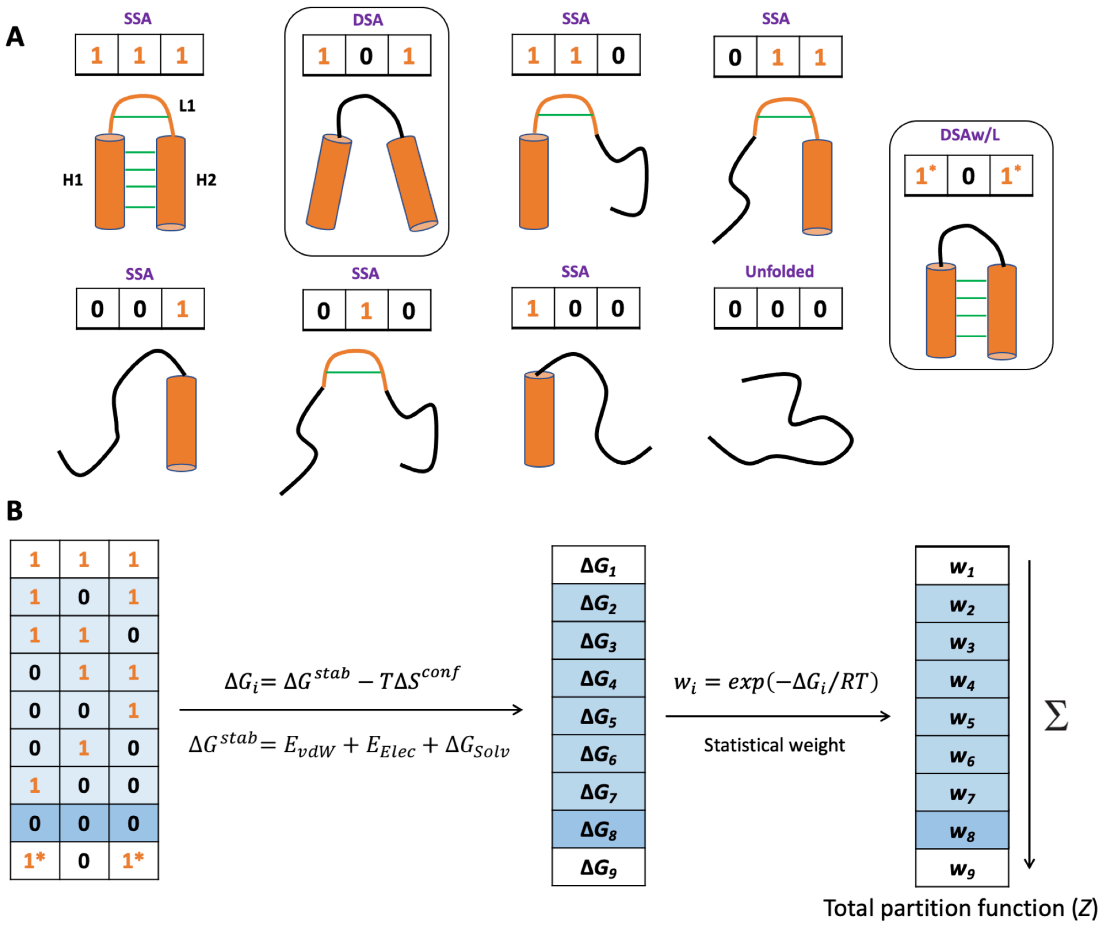


**Figure S1** Microstate definitions in the WSME model. (A) Model microstates for a highly coarse-grained version of a protein where the *entire* secondary structure elements (H1, L1, and H2) are considered as blocks. *1* represents the folded block and *0* represented the unfolded block. For such a three-block representation, there can be a maximum of 6 SSA states (single sequence approximation), one defined by DSA (double sequence approximation), one DSAw/L state (DSA with loop) and one fully unfolded state. In DSA (boxed and adjacent to the fully folded protein), the helices H1 and H2 do not interact as L1 is unfolded, while in DSAw/L the two helices can interact despite the unfolded L1 (the state to the extreme right of the panel; asterisks are employed to distinguish the state from DSA). The interactions within the secondary structure elements are also considered. *In the current work, we employ a similar formalism for GPCRs but coarse-grain the protein in four-residue blocks, resulting in millions of such conformational substates (Table S1).* (B) Since the structure of the each of the microstates are known (from the PDB file), the corresponding free-energy of microstates ($\Delta G_i$) can be calculated considering the balance between stabilization free energy ($\Delta G^{stab}$) and conformational entropy ($\Delta S^{conf}$). The former includes contributions from van der Waals interactions (vdW), electrostatics (Elec) and solvation (empirical solvation free-energy term; see main text). From the free energies, the statistical weights and hence the total partition function can be calculated (summation of statistical weights). The partition function can, in turn, be employed to calculate heat capacity profiles, free-energy profiles and free-energy landscapes (see main text).

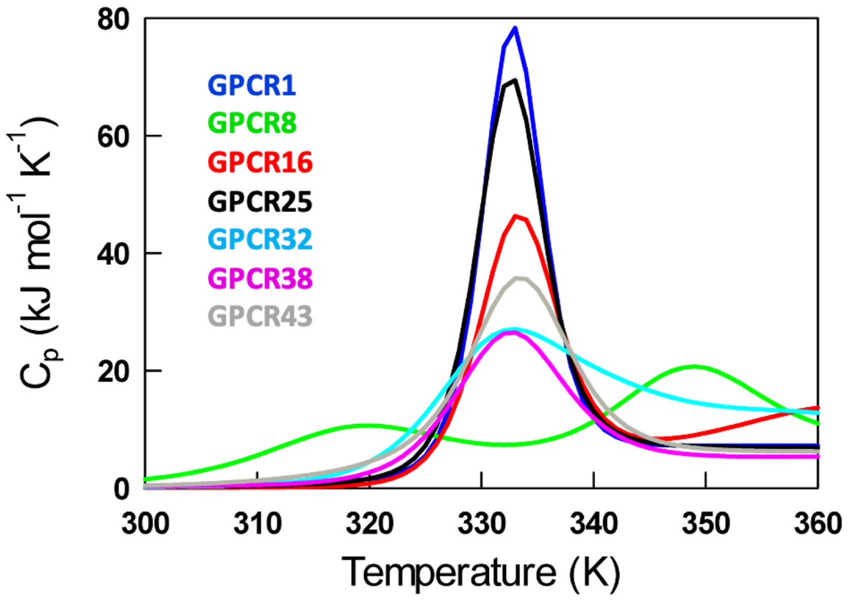


**Figure S2** Predicted heat capacity profiles of representative GPCRs that span the spectrum of thermodynamic unfolding behaviors. See Table S1 for the name and PDB ids of GPCRs.

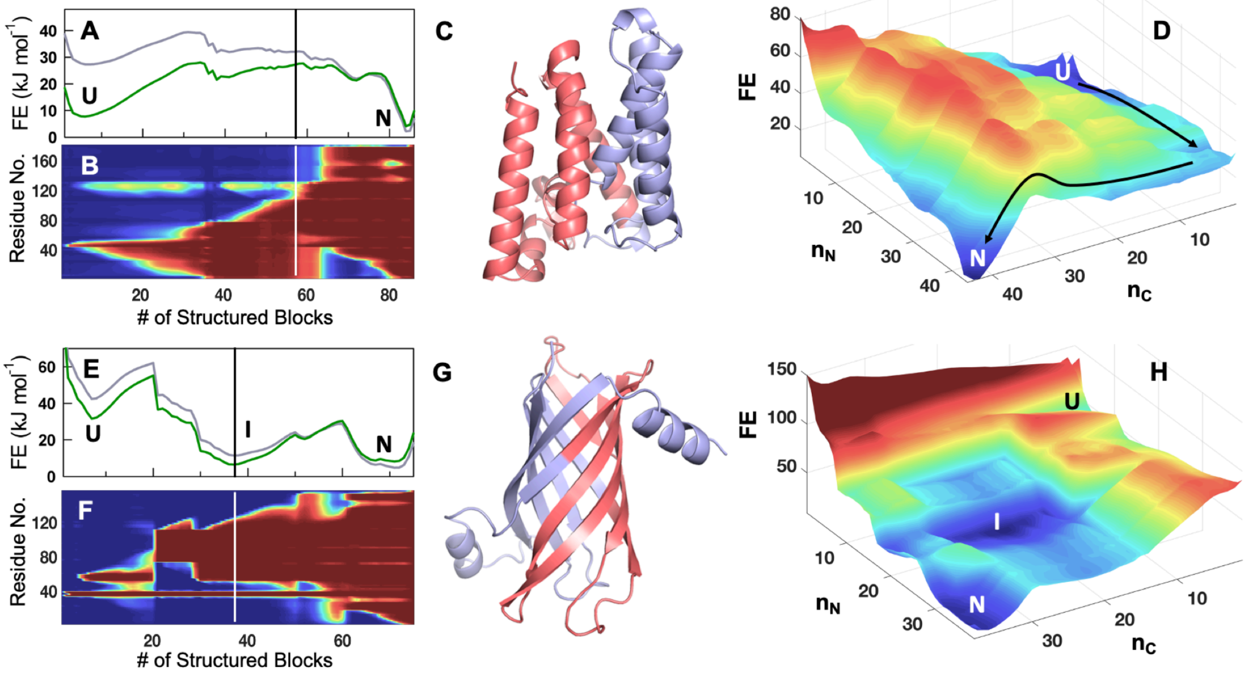


**Figure S3** (A-D) Conformational landscape of the helical membrane protein GlpG. Panel A plots the free energy profile at 310 K and the melting temperature of 333 K (grey and green, respectively) as a function of the reaction coordinate, the number of structured blocks. Panel B plots the folding probability of different protein regions (y-axis) as a function of the reaction coordinate (x-axis) in the spectral scale (blue represents zero folding probability while red represents a folding probability of one). At 57 structured blocks (the vertical dashed in panels A and B), the N-terminal half is more folded than the C-terminal half. Protein regions that are folded and unfolded at 57 structured blocks are mapped onto the structure in salmon and light blue, respectively. A similar observation on the folding mechanism can be made from the perspective of the two-dimensional folding landscape as a function of the number of folded structured blocks in the N- and C-terminal halves ($n_N$ and $n_C$, respectively). The vertical axis is the free energy (FE) in kJ mol$^{-1}$. The arrows point to the direction of folding starting from the unfolded state (U) to the folded state (N). It can be seen that the N-terminal region folds first followed by the C-terminal region. Note that these inferences are made from equilibrium populations and not from kinetic modeling that can potentially reveal higher complexity. (E-H) Folding mechanism of the beta-sheet-rich PagP following the same color code as panels A-D. In panel F, it can be seen that the C-terminal half is relatively more structured compared to the N-terminal half; this is mapped onto the structure in panel G.  In PagP, the salmon colored region in panel G forms an intermediate (I). The intermediate can also be observed in the 2D landscape.

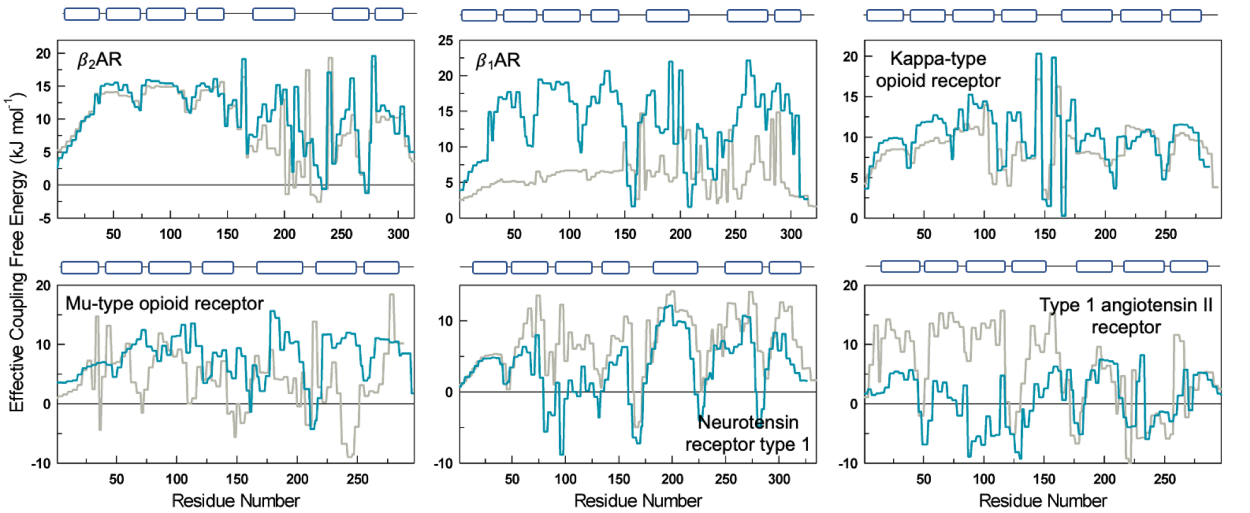


**Figure S4** Effective coupling energies for the inactive (gray) and active (dark cyan) forms of the GPCRs. The cartoon on top displays the length and position of the transmembrane helices. Also refer to Figure 6 in the main text and the associated discussion.

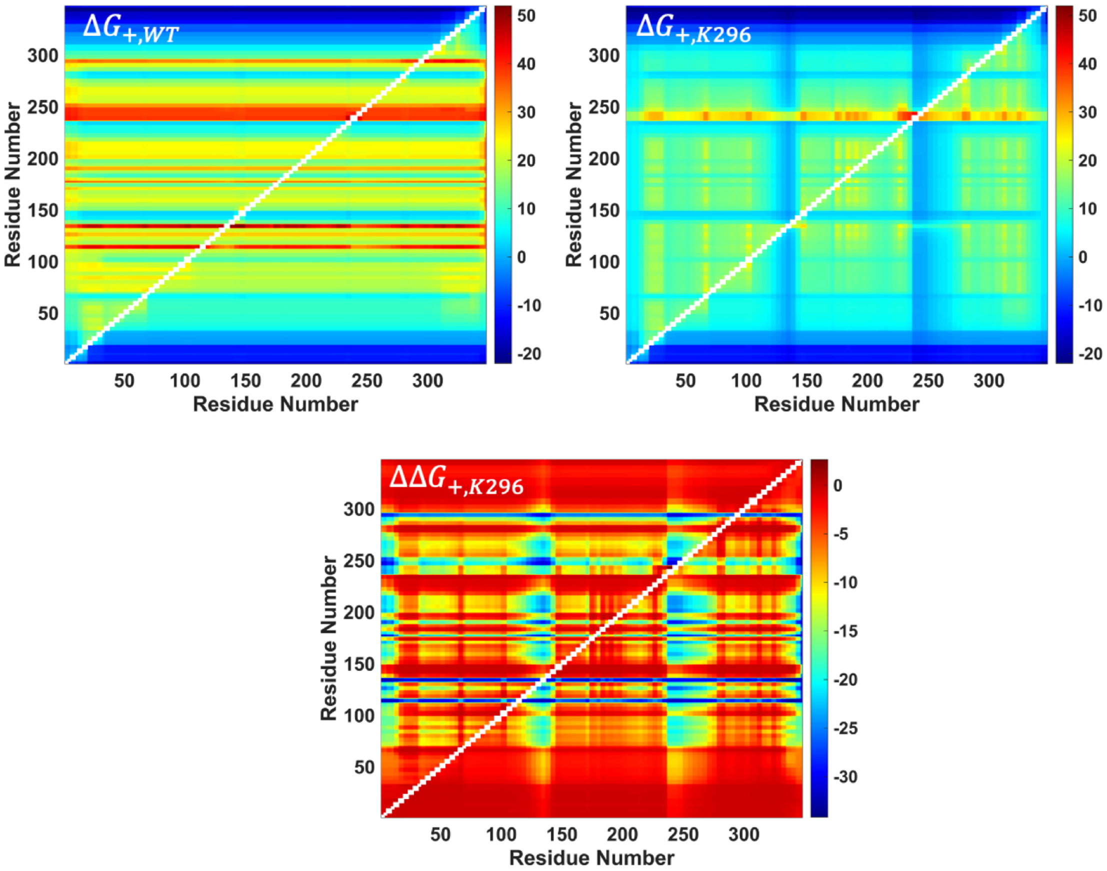


**Figure S5** The positive coupling matrix for WT Rhodopsin (top left), the mutant K296 (top right) and the difference between the two (bottom row). It can be seen that the residues in the mutant are less coupled than the WT (top right).

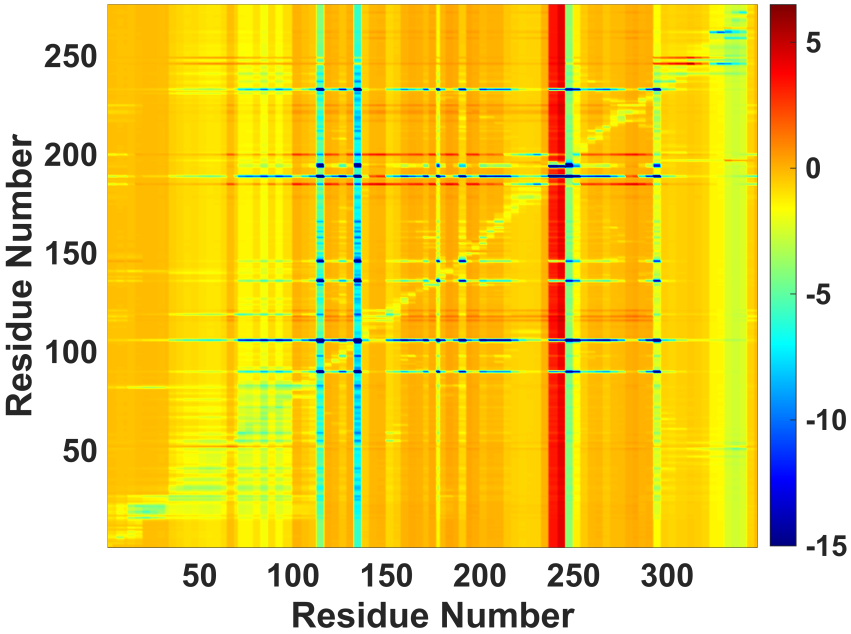


**Figure S6** Mutational response matrix - every row in this matrix represents the $\langle \Delta\Delta G_{+} \rangle$ across all residues for a specific alanine mutation or perturbation in Rhodopsin. The average and standard deviation across the rows is shown as Figures 7B and 7C in the main text. The lowest $\langle \Delta\Delta G_{+} \rangle$ is -47 kJ mol$^{-1}$, but the color range at the lower end of the spectrum is constrained here to -15 kJ mol$^{-1}$ for the sake of clarity.

**Table S1.** Database of inactive GPCR structures.

| GPCR Index | Protein | PDB ID | UniProt ID | Class | No. of Residues (*N*) | No. of Blocks ($N_b$) | No. of Microstates |
|---|---|---|---|---|---|---|---|
| 1 | Rhodopsin | 1U19 | P02699 | A | 349 | 90 | 5,349,436 |
| 2 | C-X-C chemokine receptor type 1 | 2LNL | P25024 | A | 296 | 77 | 2,855,854 |
| 3 | Beta-2 adrenergic receptor | 2RH1 | P07550 | A | 314 | 84 | 4,053,141 |
| 4 | C-X-C chemokine receptor type 4 | 3ODU | P61073 | A | 293 | 75 | 2,568,801 |
| 5 | Sphingosine 1-phosphate receptor 1 | 3V2Y | P21453 | A | 311 | 82 | 3,678,644 |
| 6 | Proteinase-activated receptor 1 | 3VW7 | P25116 | A | 288 | 77 | 2,855,854 |
| 7 | Beta-1 adrenergic receptor (Turkey) | 4BVN | P07700 | A | 323 | 86 | 4,455,532 |
| 8 | Kappa-type opioid receptor | 4DJH | P41145 | A | 293 | 76 | 2,709,477 |
| 9 | Mu-type opioid receptor (Mouse) | 4DKL | P42866 | A | 288 | 76 | 2,709,477 |
| 10 | Delta-type opioid receptor | 4N6H | P41143 | A | 303 | 78 | 3,008,084 |
| 11 | Metabotropic glutamate receptor 1 | 4OR2 | Q13255 | C | 263 | 69 | 1,836,206 |
| 12 | P2Y purinoceptor 12 | 4PXZ | Q9H244 | A | 291 | 75 | 2,568,801 |
| 13 | Neurotensin receptor type 1 | 4XES | P20789 | A | 335 | 90 | 5,349,436 |
| 14 | P2Y purinoceptor 1 | 4XNV | P47900 | A | 296 | 76 | 2,709,477 |
| 15 | G-protein coupled receptor homolog US28 [Human cytomegalovirus (strain AD169) (HHV-5) (Human herpesvirus 5)] | 4XT1 | P69332 | Viral | 296 | 79 | 3,166,321 |
| 16 | Type-1 angiotensin II receptor | 4YAY | P30556 | A | 306 | 79 | 3,166,321 |
| 17 | Lysophosphatidic acid receptor 1 | 4Z35 | Q92633 | A | 304 | 81 | 3,501,442 |
| 18 | Orexin receptor type 1 | 4ZJ8 | O43613 | A | 348 | 92 | 5,843,749 |
| 19 | Nociceptin receptor | 5DHG | P41146 | A | 285 | 74 | 2,433,676 |

| | | | | | | | |
|---|---|---|---|---|---|---|---|
| 20 | C-C chemokine receptor type 9 | 5LWE | P51686 | A | 298 | 76 | 2,709,477 |
| 21 | Proteinase-activated receptor 2 | 5NDD | P55085 | A | 301 | 78 | 3,008,084 |
| 22 | Free fatty acid receptor 1 | 5TZR | O14842 | A | 283 | 73 | 2,303,954 |
| 23 | Adenosine receptor A1 | 5UEN | P30542 | A | 308 | 82 | 3,678,644 |
| 24 | C-C chemokine receptor type 5 | 5UIW | P51681 | A | 301 | 79 | 3,166,321 |
| 25 | Type-2 angiotensin II receptor | 5UNF | P50052 | A | 292 | 74 | 2,433,676 |
| 26 | Apelin receptor | 5VBL | P35414 | A | 312 | 78 | 3,008,084 |
| 27 | Glucagon-like peptide 1 receptor | 5VEW | P43220 | | 287 | 77 | 2,855,854 |
| 28 | Neuropeptide Y receptor type 1 | 5ZBQ | P25929 | A | 320 | 85 | 4,250,766 |
| 29 | Platelet-activating factor receptor | 5ZKP | P25105 | A | 310 | 80 | 3,330,721 |
| 30 | Cannabinoid receptor 2 | 5ZTY | P34972 | A | 299 | 76 | 2,709,477 |
| 31 | 5-hydroxytryptamine receptor 2A | 6A94 | P28223 | A | 330 | 87 | 4,667,609 |
| 32 | Frizzled-4 | 6BD4 | Q9ULV1 | F | 333 | 89 | 5,114,386 |
| 33 | C5a anaphylatoxin chemotactic receptor 1 | 6C1R | P21730 | A | 298 | 78 | 3,008,084 |
| 34 | Prostaglandin D2 receptor 2 | 6D27 | Q9Y5Y4 | A | 323 | 83 | 3,862,489 |
| 35 | Metabotropic glutamate receptor 5 | 6FFI | P41594 | C | 264 | 68 | 1,731,349 |
| 36 | C-C chemokine receptor type 2 | 6GPX | P41597 | A | 286 | 75 | 2,568,801 |
| 37 | Substance-P receptor | 6HLP | P25103 | A | 301 | 77 | 2,855,854 |
| 38 | Endothelin receptor type B | 6IGK | P24530 | A | 318 | 82 | 3,678,644 |
| 39 | Thromboxane A2 receptor | 6IIU | P21731 | A | 314 | 84 | 4,053,141 |
| 40 | Prostaglandin E2 receptor EP3 subtype | 6M9T | P43115 | A | 308 | 82 | 3,678,644 |
| 41 | Melatonin receptor type 1A | 6ME2 | P48039 | A | 299 | 79 | 3,166,321 |
| 42 | Melatonin receptor type 1B | 6ME6 | P49286 | A | 294 | 76 | 2,709,477 |

| | | | | | | | |
|---|---|---|---|---|---|---|---|
| 43 | Calcitonin receptor | 6NIY | P30988 | B | 277 | 74 | 2,433,676 |
| 44 | C-C chemokine receptor type 7 | 6QZH | P32248 | A | 290 | 73 | 2,303,954 |
| 45 | Cysteinyl leukotriene receptor 2 | 6RZ6 | Q9NS75 | A | 294 | 75 | 2,568,801 |

**Table S2.** Model parameters and thermodynamic outputs.

| GPCR Index | Protein | PDB ID | vdW Interaction Energy (J mol$^{-1}$) | Strongly Coupled Residues at 310 K ($f_c$ in %) | No. of Intermediates from 1D FE Profile at $T_m$ (1*RT* threshold) |
|---|---|---|---|---|---|
| 1 | Rhodopsin | 1U19 | -48.2 | 9.8 | 1 |
| 2 | C-X-C chemokine receptor type 1 | 2LNL | -56.3 | 9.4 | 0 |
| 3 | Beta-2 adrenergic receptor | 2RH1 | -46.7 | 15.6 | 2 |
| 4 | C-X-C chemokine receptor type 4 | 3ODU | -52.0 | 23.9 | 0 |
| 5 | Sphingosine 1-phosphate receptor 1 | 3V2Y | -49.2 | 7.4 | 3 |
| 6 | Proteinase-activated receptor 1 | 3VW7 | -52.3 | 16.7 | 0 |
| 7 | Beta-1 adrenergic receptor (Turkey) | 4BVN | -46.7 | 11.1 | 1 |
| 8 | Kappa-type opioid receptor | 4DJH | -47.4 | 7.5 | 2 |
| 9 | Mu-type opioid receptor (Mouse) | 4DKL | -49.9 | 12.2 | 0 |
| 10 | Delta-type opioid receptor | 4N6H | -46.1 | 19.1 | 2 |
| 11 | Metabotropic glutamate receptor 1 | 4OR2 | -48.6 | 10.6 | 1 |
| 12 | P2Y purinoceptor 12 | 4PXZ | -47.6 | 21.0 | 2 |
| 13 | Neurotensin receptor type 1 | 4XES | -53.2 | 20.3 | 1 |
| 14 | P2Y purinoceptor 1 | 4XNV | -46.6 | 12.2 | 6 |
| 15 | G-protein coupled receptor homolog US28 [Human cytomegalovirus (strain AD169) (HHV-5) (Human herpesvirus 5)] | 4XT1 | -48.3 | 7.1 | 3 |
| 16 | Type-1 angiotensin II receptor | 4YAY | -50.1 | 21.6 | 0 |
| 17 | Lysophosphatidic acid receptor | 4Z35 | -51.9 | 8.9 | 0 |

| | | | | | |
|---|---|---|---|---|---|
| | 1 | | | | |
| 18 | Orexin receptor type 1 | 4ZJ8 | -49.70 | 10.3 | 6 |
| 19 | Nociceptin receptor | 5DHG | -54.4 | 8.4 | 2 |
| 20 | C-C chemokine receptor type 9 | 5LWE | -45.2 | 15.1 | 1 |
| 21 | Proteinase-activated receptor 2 | 5NDD | -46.0 | 13.0 | 0 |
| 22 | Free fatty acid receptor 1 | 5TZR | -52.4 | 15.2 | 0 |
| 23 | Adenosine receptor A1 | 5UEN | -47.3 | 10.1 | 2 |
| 24 | C-C chemokine receptor type 5 | 5UIW | -45.6 | 9.6 | 0 |
| 25 | Type-2 angiotensin II receptor | 5UNF | -50.2 | 11.3 | 2 |
| 26 | Apelin receptor | 5VBL | -50.2 | 15.7 | 1 |
| 27 | Glucagon-like peptide 1 receptor | 5VEW | -47.9 | 13.2 | 4 |
| 28 | Neuropeptide Y receptor type 1 | 5ZBQ | -48.5 | 20.0 | 1 |
| 29 | Platelet-activating factor receptor | 5ZKP | -48.3 | 18.4 | 2 |
| 30 | Cannabinoid receptor 2 | 5ZTY | -47.3 | 14.7 | 1 |
| 31 | 5-hydroxytryptamine receptor 2A | 6A94 | -48.2 | 16.1 | 0 |
| 32 | Frizzled-4 | 6BD4 | -45.7 | 9.6 | 1 |
| 33 | C5a anaphylatoxin chemotactic receptor 1 | 6C1R | -47.8 | 7.0 | 4 |
| 34 | Prostaglandin D2 receptor 2 | 6D27 | -52.3 | 16.7 | 3 |
| 35 | Metabotropic glutamate receptor 5 | 6FFI | -49.8 | 15.2 | 0 |
| 36 | C-C chemokine receptor type 2 | 6GPX | -47.7 | 16.8 | 2 |
| 37 | Substance-P receptor | 6HLP | -48.7 | 12.3 | 3 |
| 38 | Endothelin receptor type B | 6IGK | -45.2 | 11.3 | 1 |
| 39 | Thromboxane A2 receptor | 6IIU | -49.3 | 10.2 | 3 |

| | | | | | |
|---|---|---|---|---|---|
| 40 | Prostaglandin E2 receptor EP3 subtype | 6M9T | -56.0 | 3.9 | 3 |
| 41 | Melatonin receptor type 1A | 6ME2 | -48.1 | 9.4 | 1 |
| 42 | Melatonin receptor type 1B | 6ME6 | -46.8 | 9.5 | 1 |
| 43 | Calcitonin receptor | 6NIY | -48.0 | 13.0 | 1 |
| 44 | C-C chemokine receptor type 7 | 6QZH | -45.7 | 12.8 | 5 |
| 45 | Cysteinyl leukotriene receptor 2 | 6RZ6 | -47.8 | 10.9 | 0 |

**Table S3.** Model parameters for GlpG and PagP.

| Protein | GlpG | PagP |
|---|---|---|
| **PDB ID** | 2XOV | 1THQ |
| **No. of Residues** | 181 | 157 |
| **Block size** | 2 | 2 |
| **No. of Blocks** | 86 | 75 |
| **No. of Microstates** | 4,455,532 | 2,568,801 |
| **vdW Interaction Energy (J $mol^{-1}$)** | -49.0 | -46.5 |

**Table S4.** Model parameters employed for predicting free energy profiles from active and inactive structures at 310 K. (note that the parameters for inactive structures are same in Table S2)

| GPCR Index | Inactive Structures | | | Active Structures | | |
|---|---|---|---|---|---|---|
| | PDB ID | Resolution (Å) | vdW Interaction Energy (J $mol^{-1}$) | PDB ID | Resolution (Å) | vdW Interaction Energy (J $mol^{-1}$) |
| 1 | 1U19 | 2.2 | -48.2 | 5W0P | 3.0 | -47.8 |
| 3 | 2RH1 | 2.4 | -46.7 | 4LDE | 2.8 | -48.7 |
| 7 | 4BVN | 2.1 | -46.7 | 6H7N | 2.5 | -50.3 |
| 8 | 4DJH | 2.9 | -47.4 | 6B73 | 3.1 | -49.5 |
| 9 | 4DKL | 2.8 | -49.9 | 5C1M | 2.1 | -49.6 |
| 13 | 4XES | 2.6 | -53.2 | 6OS9 | 3.0 | -55.0 |
| 16 | 4YAY | 2.9 | -50.1 | 6DO1 | 2.9 | -53.1 |
| 23 | 5UEN | 3.2 | -47.3 | 6D9H | 3.6 | -48.9 |
